# Performance efficient macromolecular mechanics via sub-nanometer shape based coarse graining

**DOI:** 10.1101/2022.08.28.505590

**Authors:** Alexander J. Bryer, Juan R. Perilla

**Affiliations:** Department of Chemistry and Biochemistry, University of Delaware, Newark, DE, 19716

## Abstract

Dimensionality reduction via coarse grain modeling has positioned itself as an indispensable tool for decades, particularly for biomolecular simulations where atomic systems encompass hundreds of millions of atoms. While distinct flavors of coarse grain modeling exist, those occupying the coarse end of the spectrum are typically knowledge based, relying on a priori information to parameterize models, thus hindering general predictive capability. Here, we present an algorithmic and transferable approach known as shape based coarse graining (SBCG) which employs unsupervised machine learning via competitive Hebbian adaptation to construct coarse molecules that perfectly represent atomistic topologies. We show how SBCG provides ample control over model granularity, and we provide a quantitative metric for selection thereof. Parameter optimization, inclusion of small molecule species, as well as simulation configuration are discussed in detail. Our method and its implementation is made available as part of the CGBuilder plugin, present in the widely-used visual molecular dynamics (VMD) and nanoscale molecular dynamics (NAMD) software suites. We demonstrate applications of our method with a variety of systems from the inositol hexaphosphate-bound, full-scale HIV-1 capsid to heteromultimeric cofilin-2-bound actin filaments. Overall, we show that SBCG provides a simple yet robust approach to coarse graining that requires minimal user input and lacks any ad hoc interactions between protein domains. Furthermore, because the Hamiltonian employed in SBCG is CHARMM compatible, SBCG takes full advantage of the latest GPU-accelerated NAMD3 yielding molecular sampling of over a microsecond per day for systems that span micrometers.

## Main

Molecular dynamics (MD) simulations evolve chemical systems over time via integration of Newton’s equations of motion [1]. Since its inception as an investigatory method, MD simulation has provided high spatial and temporal resolution data of materials, surfaces and biomolecular systems that complement experimentally-derived information. A widely-recognized challenge in the domains of computational biochemistry and physics, which has driven the development of novel hardware [2] and software alike, is the computational complexity of biomolecular systems.

As early as 1975, dimensionally-reduced descriptions of molecular systems have been employed to lessen the computational cost associated with protein folding simulation [3]. This practice, referred to generally as coarse graining (CG), produces models that seek to accurately represent chemical systems with far fewer degrees of freedom than are present at atomistic resolution (3*N* − 1). The scope and strategy of CG simulations have been revolutionized many times over in the last ∼50 years and CG modeling has been successfully applied to gas, liquid and condensed phase systems [4, 5, 6, 7, 8]. While the geometric increase in computing power of the late 20th century until present has made atomistic simulation more computationally tractable, CG modeling has remained a staple of computational science. Considering our growing understanding of climate change and the extreme energy costs of supercomputing, the latter which continues to balloon with ever-increasing computing power, we anticipate that dimensionality reduction via CG modeling will remain a staple for decades to come.

In general, coarse graining refers to mapping, by various criteria, groups of atoms in ℝ^3^ to a single position, or bead. In the present context, the term *granularity* refers to the degree of reduction, i.e., the coarseness of a given model, or how many atoms are mapped to a single bead. Granularity depends on several factors and is, in general, established by the scientist based on the nature of their system and the questions they seek to investigate.

On the high-granularity end of the spectrum, MARTINI [9, 10, 11] is a popular offering. Parameterized empirically, MARTINI maps four atoms to a single bead. The MARTINI force field contains bond, angle, dihedral and non-bonded interaction terms that govern model behavior. On the low-granularity end of the spectrum, so-called ultra coarse grained (UCG) models map many atoms, sometimes entire protein domains in biomolecular simulation, to a single bead [12]. Particularly adept at modeling large assemblies [13] as well as processes of self-assembly [14, 15, 16], UCG simulations also commonly employ large integration time steps [17] that increase sampling efficiency and aid the resolution of long timescale behavior. The flexibility and efficiency of low granularity CG has enabled the study of flagellar motility [18], as well as large scale DNA dynamics via alternate levels of theory such as worm-like chain modeling [19]. Resulting from significant information loss due to extreme coarseness, low granularity, UCG models are often knowledge-based; that is, the interactions among constituent beads are often established explicitly to reproduce known behavior in the absence of charge, hydrophobicity or other physical properties lost in reduction. Multiscale CG models [20] and force fields, e.g., SIRAH [21], PACE [22, 23] and others, span the gap between low- and high-granularity regimes and have proven especially useful in CG descriptions of aqueous environments.

Construction and parameterization of CG models of varying granularity has also motivated the usage of machine learning [24, 25, 26, 27, 28]. The high dimensionality of molecular models, even coarse models, lends utility to neural networks suited for highly dimensional optimization problems. The general paradigm of machine learning is to evolve a set of numerical weights, given a particular input pattern, in response to an objective function. Commonly, the optimization is supervised. That is, the objective function is supplied with training data, often experimental or high-resolution data, as e.g. collected from atomistic simulation, to evaluate and evolve network parameters. This flexible approach has been successfully utilized in a variety of settings, including generalized and systematic CG force field optimization [24] and CG modeling of water [25]. Derivation of CG force fields for dissipative particle dynamics (DPD) has also been accomplished with a Bayesian network [27], another supervised scheme employing Bayesian inference for parameter optimization in the absence of a fully-realized objective function. A specific class of models, generative adversarial networks (GANs), employ competing prediction networks in a zero-sum game and learn to reproduce training data through optimization. GANs represent a semi- or weakly-supervised paradigm and have been successfully applied to derivation of CG models [26]. Another hybrid architecture permitting weak supervision is the graph neural network (GNN), and this has also been productively utilized to optimize CG force fields [28]. Due to the dependence of supervised learning on robust, unbiased and voluminous training data, unsupervised learning is an attractive strategy for a variety of problems.

A separate class of analytical methods for deriving CG potentials from atomistic data has also been developed. These frameworks, built on numerical optimization, have found significant utility at large scales. Approaches such as Lennard-Jones (LJ) static potential matching employ numerical optimization to define the LJ interaction potential of CG beads based on an atomistic force field [29]. Other techniques such as force matching and Boltzmann inversion have been successfully utilized to derive force field terms in numerous CG contexts, ranging from organic polymers [30] to meso and multiscale biomolecular assemblies [31, 20, 32, 30]. The relative entropy approach is an alternative optimization method that is additionally able to quantify the error of a CG model relative to a model built from first principles [33]. Similar to unsupervised learning methods, analytical approaches to deriving CG potentials have low dependence on high resolution data compared to supervised learning via, e.g., a convolutional neural network. These methods are well-suited for the efficient parameterization of macromolecular complexes.

The criteria that a CG reduction employs to yield models with particular granularities are often defined *ad hoc*. For example, a scientist who aims to simulate a viral capsid, encompassing tens of millions of atoms, may elect to employ a UCG model, and she might establish only one or two CG beads per capsid subunit based on information such as the presence of independently-folded protein domains. Conversely, the desired timescale of the model may necessitate an UCG description to access larger integration time steps. From experimental or high-resolution computational data, such as principal component analysis of an atomistic counterpart, the model is Parameterized. For instance, parameterization of coarse models such as Elastic Network Models (ENMs) might consider and attempt to reproduce through a bonded network relative motions and essential dynamics of subunits in the capsid assembly [34, 35]. While serviceable to the foundational goals of the effort, such models suffer from model dependent realism, and thus their predictive capabilities are contextually limited.

A desirable quality of CG modeling is *transferability*, i.e., readiness of the CG reduction to be applied to other structures and systems while retaining general predictive potential [36, 29]. To satisfy this requirement, the CG reduction should be algorithmic and thus predictable. Further, the reduction should retain enough information of the atomistic input to faithfully reproduce its natural properties, such as multimeric assembly characteristics, without the need for explicit parameterization.

Herein, we will present arguments and evidence that shape-based coarse graining (SBCG) is a highly flexible and transferable CG flavor that satisfies such requirements, and is markedly successful in modeling biomolecular assemblies. The driving force of SBCG is a Topology Representing Network (TRN), which employs competitive Hebbian adaptation as part of an unsupervised learning framework to perform 3-D Voronoi tessellation and generate perfectly topology preserving CG molecules. We will describe changes to the original algorithm that enable SBCG mappings entering the high-granularity regime, and show that resulting models conform to atomistic charge density profiles within sub-nanometer resolution. Next, we will describe parameter optimization via iterative Boltzmann inversion and discuss considerations for its deployment in high-granularity, sub-nanometer use-cases. We demonstrate the utility of this method via application to three unique protein structures, comprising two macromolecular biological assemblies: human cofilin-2 bound to actin filaments [37] and the full-scale HIV-1 conical capsid. Finally, we will provide guidance for configuring and performing SBCG simulations.

## Results

To assess the transferability and performance of our SBCG modeling approach, we sought three unique molecular applications with the goal of simulating both homo- and hetero-multimeric assemblies: HIV-1 CA, assembled into a conical capsid (Figure 1), and cofilin-2 bound to actin filaments (Figure 2). Utilizing our sub-nanometer SBCG modeling approach, we constructed, validated and parameterized our macromolecular systems, then performed NVT simulations using the fully GPU-resident NAMD3 molecular dynamics engine [38].

**Figure 1:**
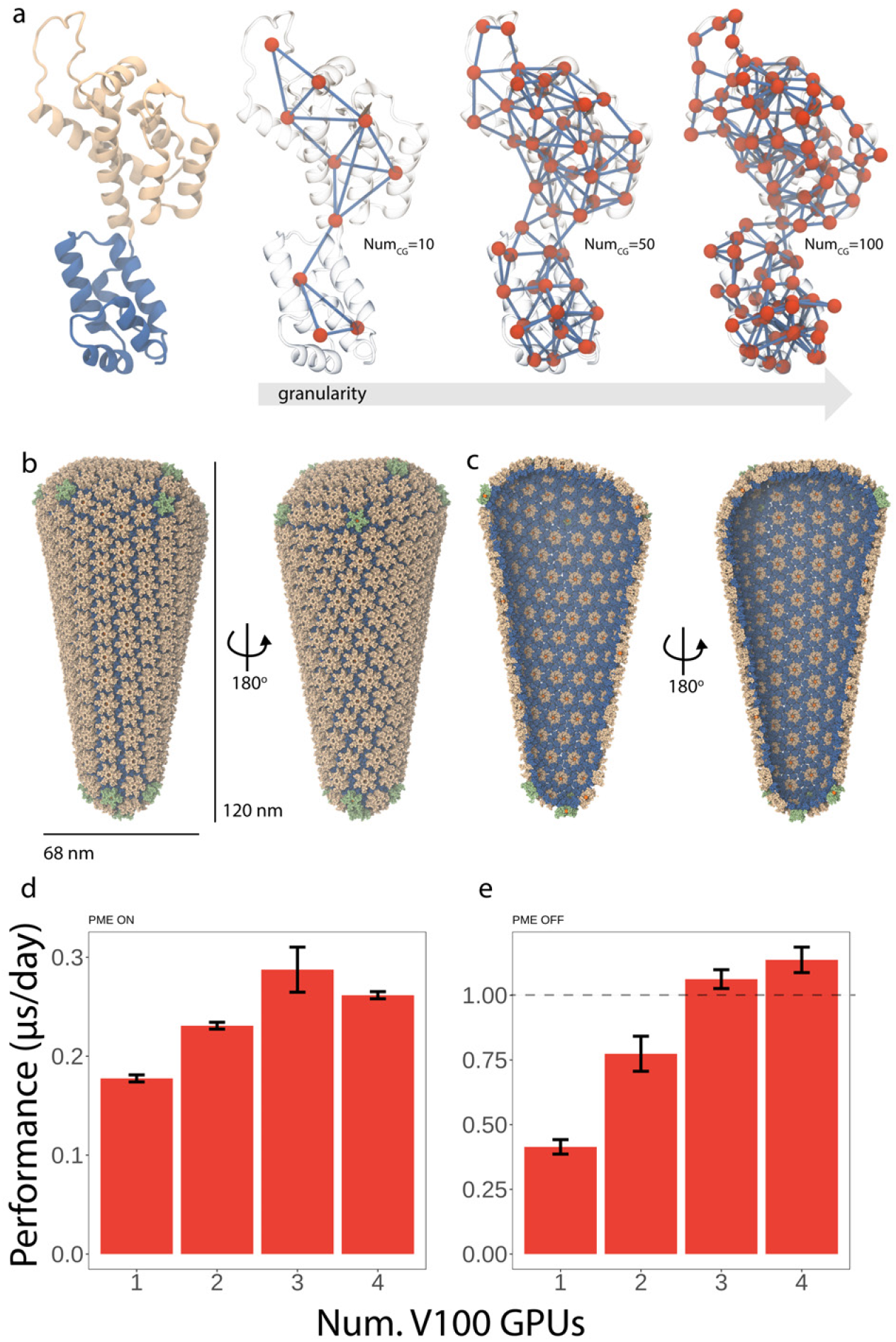
Shape based coarse grain (SBCG) HIV-1 capsid. **a** View of the HIV-1 CA monomer, left, colored by domain. On the right, three corresponding SBCG models of HIV-1 CA with increasing granularity. Granularity in this case refers to the number of beads employed to model the structure. **b** Full view of the SBCG HIV-1 conical capsid, shown from two perspectives. **c** Clipped view of the SBCG HIV-1 conical capsid, shown from two perspectives. For panels b and c, protein is shown as vdW beads, with the CA amino terminal domains colored tan and the CA carboxy terminal domains colored blue. IP6 beads are shown as orange vdW spheres. [40, 39]. **d,e** Performance benchmarks with NAMD3 [38], simulating the HIV-1 conical capsid shown in panels c and d. Benchmarks were performed with one CPU per GPU employed. For both benchmarks, identical configuration was employed, and only the usage of PME for long range electrostatics varied. The time step employed was 48 fs per step; Langevin *γ* was set to 2.0 ps^−1^; bonded interactions were evaluated every time step and nonbonded interactions were evaluated every other timestep. **d** NAMD3 GPU benchmarks, utilizing NVIDIA V100s, with PME *on*. **e** NAMD3 GPU benchmarks, utilizing NVIDIA V100s, with PME *off*. Remarkably, for three and four GPUs per simulation we exceed one microsecond per day simulation performance (dashed line).

**Figure 2:**
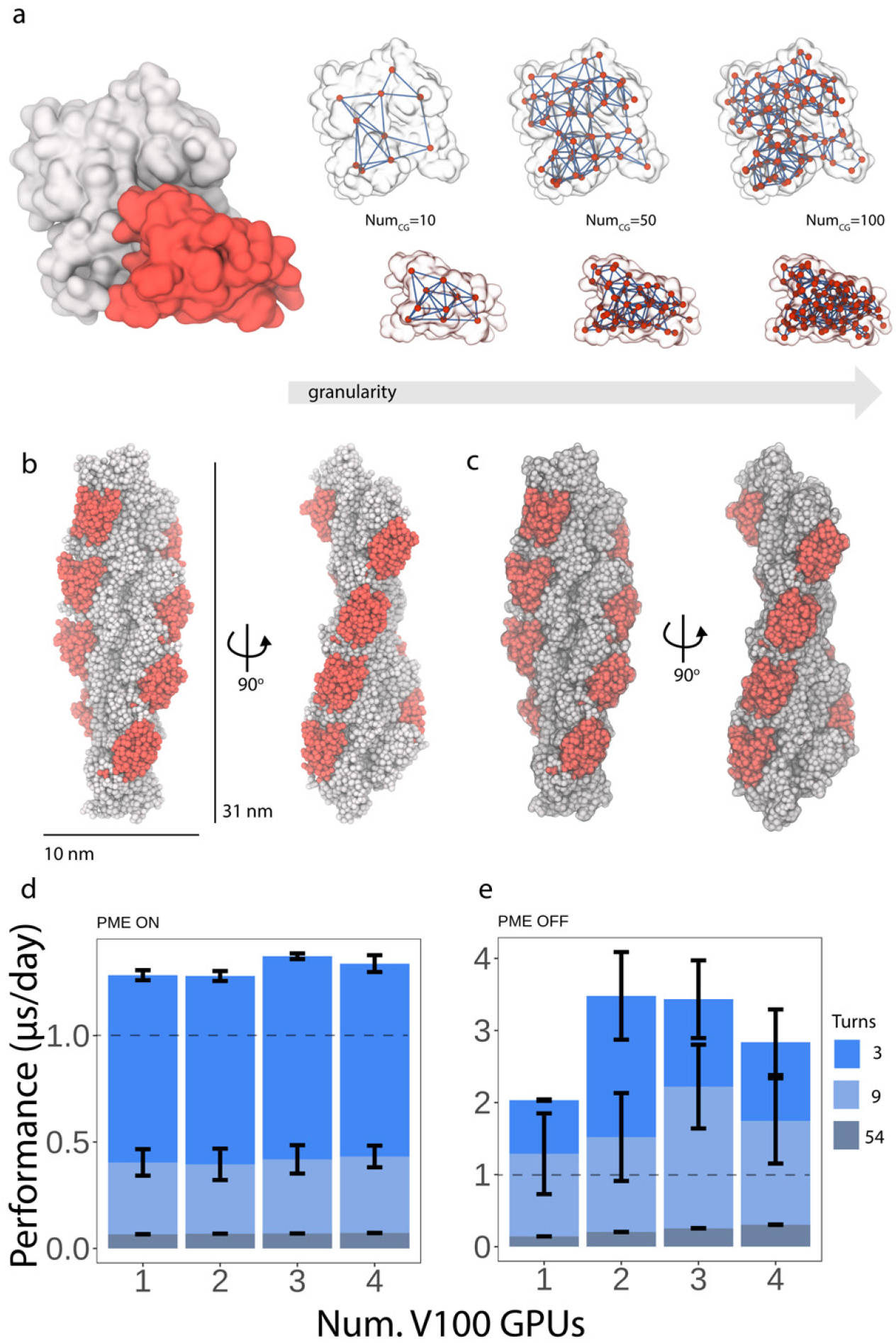
SBCG heteromultimeric cofilin-2 on actin filaments. **a** Left, atomistic surface representation of globular actin (white) bound to cofilin-2 (red). Right, corresponding SBCG representations of actin and cofilin-2, shown superimposed with the atomistic molecular surfaces. **b** A single turn of the SBCG cofilin-2-bound actin filament, shown from two perspectives. One turn corresponds to a length of 31 nm. **c** The SBCG cofilin-2-bound actin filament superimposed with the atomistic molecular surface, shown transparently. The shape of the atomistic filament is perfectly represented by the coarse model. **d** NAMD3 GPU benchmarks, utilizing NVIDIA V100s with PME *on*. **e** NAMD3 GPU benchmarks, utilizing NVIDIA V100s, with PME *off*. For panels d and e, we performed benchmark simulations with 3-, 9- and 54-turn filaments to assess load-balancing and scaling with respect to system size. A legend is provided on the right.

After applying our framework to HIV-1 CA (Figure 1a), we assembled the full-scale conical capsid bound to the charged assembly co-factor inositol hexakisphosphate [39, 40] (Figure 1b,c, Figure S3). The complete SBCG system of the conical capsid is described by 340,000 beads, representing a larger than 200-fold reduction in particles compared to the atomistic conical capsid of the same morphology. Importantly, SBCG molecular dynamics simulations achieved greater than 1 microsecond per day sampling performance without PME-based electrostatic evaluation, and with PME enabled we maintain nearly 300 nanoseconds per day (Figure 1d,e) performance. For atomistic HIV-1 capsids, the capsid protein assembly equilibrates and reaches a stable configuration on the order of hundreds of nanoseconds [41]. Representing the ideal memory footprint for GPU offloading, the performance of our SBCG conical capsid system scales across multiple NVIDIA V100 GPUs and greatly broadens temporal resolution compared with traditional molecular sampling.

Our SBCG models of cofilin-2 bound to actin filaments represent significantly different assemblies from the conical capsid, at the physical and biochemical level but also at the computational level. Spatial de-composition is an important aspect of parallel molecular dynamics simulation, namely in the evaluation of nonbonded electrostatics, where the spatial domain is discretized over multiple processing entities. After employing our framework to actin and cofilin-2 (Figure 2a), we built heteromultimeric cofilin-2-bound actin filaments (Figure 2b,c) of varying length ranging from a single turn (31 nm, 7,000 beads) to a filament 54 turns in length (1.6 *μ*m, 400,000 beads). Despite the spatial challenge presented by this system, where the ratio of length to cross-sectional area is extremely large, we consistently exceeded 1 microsecond per day simulation performance with our 3-turn filament system with PME-based electrostatic evaluation (Figure 2d), but with no scaling regardless of filament size. Forgoing PME, we not only approach 4 microseconds per day performance but we once again observe scaling across multiple GPUs (Figure 2e). Optimizing the domain decomposition for PME-based evaluation is a future target of this work.

Beyond establishing performance of SBCG macromolecular assemblies, we analyzed our filamentous and conical capsid systems to gauge their stability and the efficacy of our approach. Importantly, for both systems, we specify no intermolecular, or otherwise ad hoc, interactions to maintain stability and assembly morphology. Figure 3a and b shows the 9-turn (280 nm, 69,000 beads) cofilin-2 bound actin filament at two time points. The diagram shown in Figure 3c represents the helicity of the filament at both of these time points, colored accordingly. We show that the filamentous quality, and its helical character twist and rise, are well-maintained without explicit steps to do so during model construction and optimization to enforce assembly morphology. Figure 3d shows a pairwise RMSD matrix for a 2 microsecond trajectory of the 3-turn filament at 298 K. This analysis computes the RMSD between every possible pair of structures across the whole time series, and is well-suited for comparing large assembly constructs which undergo global fluctuations. Our analysis shows that the filament reaches global stability, fluctuating within 5 Å RMSD, after approximately 200 ns.

**Figure 3:**
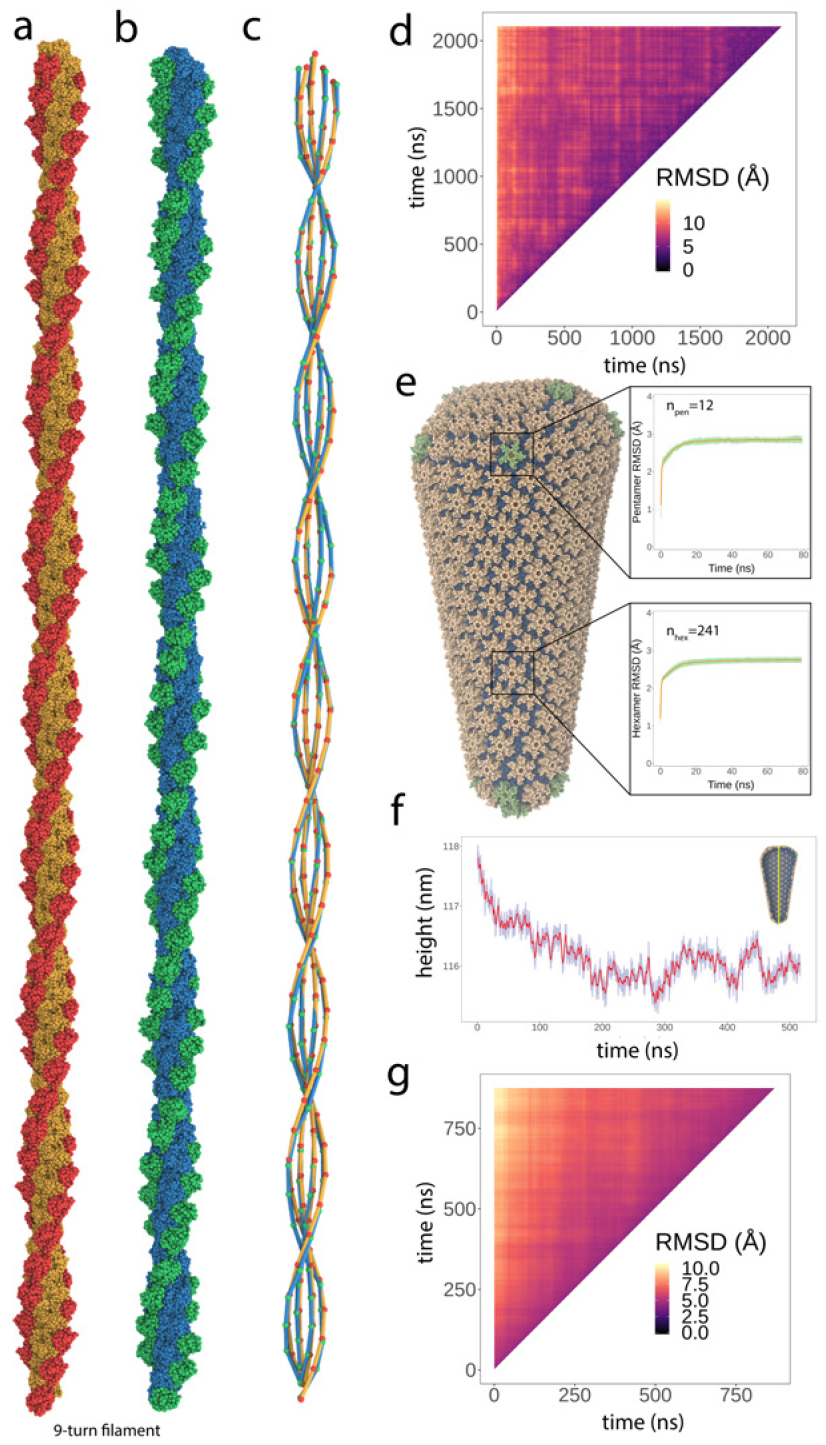
Stability of full-scale SBCG multimeric assemblies. **a**-**d** Cofilin-2 on actin filaments. **a** Initial state of the 9-turn cofilin-2-bound actin filament model, approximately 250 nm in length. **b** The 9-turn filament from panel a after unrestrained energy minimization, thermalization, equilibration and approximately 200 ns of sampling at 298 K. **c** Visualization of the 9-turn filament’s helicity computed at two time points (panels a and b). For each cofilin-2 and actin subunit comprising the assembly, the center of mass is computed and a line is drawn to its sequential neighbor along the filament’s length. The interior double helix represents actin while the outer double helix represents cofilin-2. The colors correspond to the states in panels a and b. The helical character of the filament is well-maintained throughout molecular sampling. **d** Pairwise RMSD analysis of the entire 3-turn filament, illustrated with a heatmap (colored according to the legend provided). The analysis compares every pair of structures from a 2 microsecond SBCG trajectory, yielding a matrix where every element is the RMSD between two 3-turn filaments. The analysis indicates that the filament achieves stability (< 5 Å RMSD) after roughly 200 ns. **e**-**g** HIV-1 conical capsid. **e** Pentamer and hexamer RMSD analysis from the full-scale SBCG conical capsid. For an 80 second length of trajectory yielded by equilibrium sampling, each capsomer was aligned to a single reference. The RMSD of each capsomer from the reference hexamer or pentamer was computed and the mean (orange) and standard deviation (green) was plotted across the trajectory. After approximately 20 ns of sampling at equilibrium, both hexamers and pentamers conform to the reference capsomer within 3 Å agreement. **f** Height time series of the conical capsid over 500 ns. The inset diagram illustrates the determination of height via the capsid’s principal axis of inertia. The capsid’s height converges after approximately 300 ns, and fluctuations < 1 Å are seen thereafter. **g** Pairwise RMSD analysis of the entire conical capsid, illustrated with a heatmap (colored according to the legend provided). The analysis compares every pair of structures from a 900 ns SBCG trajectory, yielding a matrix where every element is the RMSD between two complete capsids. The analysis indicates that the capsid achieves stability (< 5 Å RMSD) after roughly 300 ns.

The HIV-1 conical capsid is an irregularly shaped, closed fullerene shell. Similar to other retroviral capsids, the stability of the mature capsid manifests from intermolecular interfaces that are characterized by hydrophobicity and complementary charges. Without any ad hoc interactions to maintain stability, our SBCG capsids show marked stability. Locally, we quantify the stability of capsomers (hexamers and pentamers) via RMSD against a single reference capsomer, hexamer or pentamer, and plot the mean and standard deviation of RMSD values versus time (Figure 3e). Over approximately 20 ns of completely unrestrained sampling at 298 K, we observe capsomers achieving structural stability and converging to the reference capsomer (hexamer or pentamer) within 3 Å agreement. We employed several analyses of global behavior and stability. Figure 3f shows a trace of the capsid height over 500 ns of trajectory, computed by taking the minimum and maximum coordinates along the capsid’s principal axis of inertia. We see a small reduction in height over approximately 300 ns, then convergence and fluctuations thereafter of < 1 Å. The latter is consistent with what has been reported for a full-scale, atomistic capsid [41]. Utilizing 900 ns of sampling at 298 K, we compared every pair of frames to construct a pairwise RMSD matrix (Figure 3g). This analysis computes the RMSD between entire SBCG capsid structures, with no omissions, showing that after approximately 300 ns, the capsid achieves consistent structural agreement < 5 Å.

Our approach to SBCG utilizes the original, and only, implementation of Martinetz and Schulten’s Topology Representing Network (TRN) [42]. We introduce an exclusivity condition during initialization of neurons that enables highly granular molecular modeling. The latter, and the subsequent framework we have developed to select and validate model granularity, remove overlapping degrees of freedom and parameterize SBCG structures to match all-atom behavior, suitably models large scale macromolecules with remarkable simulation performance. We elaborate the complete SBCG modeling process, from molecular construction based on the TRN to configuring high-performance simulations, in the following sections.

## Methods

### Shape Based Coarse Graining

Shape based coarse graining (SBCG) is a modality of CG modeling which maps the coordinates of CG beads according to the shape, or topology, of an atomistic input. SBCG has been successfully employed to study the stability and deformation of viral capsids [43, 44, 45] as well as the mechanisms of lipid membrane remodeling by proteins [46, 47].

The conceptual back end of this method is a topology representing neural network [42]. The topology representing network employs a Hebbian adaptation rule with winner-take-all competition (eq. 4) to determine algorithmically the positions of CG beads relative to an atomistic input. Formally, this procedure constructs a Voronoi tessellation in ℝ^3^ [42], where each polyhedron in the tessellation represents a CG bead. The emerging Voronoi polyhedra partition atoms of the input structure, and their properties are applied to the CG bead positioned at the partition’s center-of-mass. The latter is detailed in the forthcoming sections.

#### Machine-learning based molecular topologies with competitive Hebbian adaptation

We encourage readers to refer to ref. [42] for a detailed mathematical description of the topology representing network (TRN). Here, we will first introduce basic nomenclature, then the most relevant concepts in the context of molecular topology learning, including Hebbian adaptation, Delaunay Triangulations, and finally the algorithmic formulation of the TRN itself.

For a set of neural units *i* = 1, …, *N*, lateral connections can form between any *i* to another, referred to as *j*. These lateral connections represent *synaptic links*, and are described by a matrix **C** containing connections 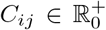. The larger an element *C_ij_* is, the stronger the synaptic link between *i* and *j*. A connection is manifest only when *C_ij_* > 0; if *C_ij_* ≤ 0 then *i* and *j* are disconnected.

Hebb’s postulate states that a pre-synaptic unit *i* shares a synaptic link with post-synaptic unit *j* if the two neural units are concurrently active. Originally formulated as a governing description of the neurological architecture of the hippocampus [50], Hebb’s rule can be represented as

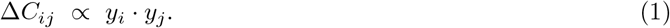

That is, the change in strength of the link between neural units *i* and *j*, Δ*C_ij_*, is proportional to the pre- and post-synaptic activity of the *i* and *j* pair. The relation in equation 1 is augmented with weight vectors 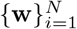 such that every neural unit *i* has a corresponding weight vector **w**_*i*_ ∈ ℝ^*D*^ which describes the center of the receptive field for neuron *i*. In this setting, the receptive field is the same as in other learning applications: it describes a region of the input space that is sensory, or responsive to stimuli, and maps to a corresponding feature or activation in the output space [51]. For constant input patterns **v** ∈ ℝ^*D*^, the activation of neural unit *i*, *y_i_*, is larger the closer its **w**_*i*_ is to **v** [42].

Martinetz and Schulten introduce a positive, continuous and monotonically decreasing function *R*(·) such that *y_i_* = *R*(||**v** − **w**_*i*_||), which describes the receptive field. This allows us to rewrite equation 1 as

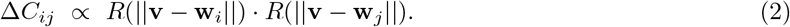

Connection strengths *C_ij_* are then solved by integrating equation 2 over the given pattern distribution *P*(**v**) as

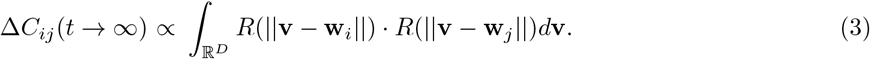

Evidently, equation 3 establishes a connection strength based simply on the area of overlap between the receptive fields of neural units *i* and *j*. Because the formulation of the receptive field *R*(·) is continuous and monotonically decreasing as ||**v − w**_*i*_|| increases, elements *C_ij_* of **C** connect all neurons to one another. The insight of Martinetz and Schulten [42] was to introduce the notion of winner-take-all selection to Hebb’s rule (eq. 1).

The competitive Hebb’s rule with winner-take-all selection becomes

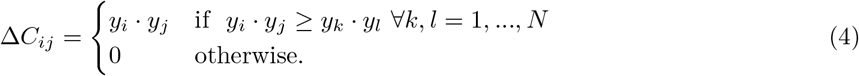

Enforcing adaptation via equation 4 rather than equation 1 results in connectivity **C** that corresponds to the Delaunay triangulation of the weight vectors **w**. Importantly, the original authors proved that, for a sequentially presented distribution of input patterns *P*(**v**) with support everywhere on ℝ^*D*^, elements *C_ij_* of **C** obey *θ*[*C_ij_*(*t* → ∞)] = *A_ij_* in the asymptotic limit. *θ*(·) is the Heavyside step function and *A_ij_* are elements of the adjacency matrix **A** of the Delaunay triangulation [42]. Here, the Delaunay triangulation is defined as the graph connecting weights **w**_*i*_ and **w**_*j*_ with adjacent Voronoi polyhedra *V_i_* and *V_j_*.

To review, elements *C_ij_* computed via the competitive Hebb’s rule (eq. 4) correspond to the adjacency matrix *A_ij_* = *θ*(*C_ij_*) for a given set of weights, points, **w** ∈ ℝ^*D*^ that denote the centers of receptive fields for all neural units *i* = 1, …, *N*. The algorithm for computing elements *C_ij_* of **C** is:

##### Algorithm 1.

(i) Initialize all connections *C_ij_* to zero;
(ii) Present input pattern **v** ∈ ℝ^*D*^ with distribution *P*(**v**);
(iii) Find unit *i* for which

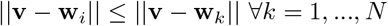

and unit *j* for which

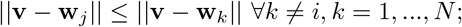
(iv) If *C_ij_* = 0, set *C_ij_* > 0 (connect *i* and *j*); else, leave *C_ij_* unchanged. Repeat at (ii).

We again invite the reader to review Theorem 1 of ref. [42] and its associated proof that *A_ij_* = *θ*(*C_ij_*) is equivalent to the adjacency matrix of the Delaunay triangulation 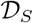 constructed from the set of weights 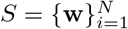.

Finally, we will introduce the topology preserving map, and particularly we will explain how competitive Hebbian adaptation as outlined above is employed for molecular topology modeling. So far, we have operated under the assumption that *P*(**v**) has support on the entire embedding space ℝ^*D*^. For many real-world input patterns, such as molecular coordinates, the input *P*(**v**) does not have support everywhere, but rather only on a submanifold *M* ⊂ ℝ^*D*^. Competitive Hebb’s rule (eq. 4) forms a subgraph of the complete Delaunay triangulation in these instances, which remains topology preserving [42].

The topology preserving map is described by a mapping Φ that projects features from a manifold *M* onto the neural units *i* = 1, …, *N* comprising a graph *G*. The mapping is directed by the set of weights 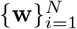 such that features of the input pattern **v** ∈ *M* are mapped to the most proximal neural unit, graph vertex, *i*. Recall that each unit *i* has an associated weight **w**_*i*_ describing its receptive field. The notation *i*\*(**v**) clarifies that the resulting Voronoi polyhedron *V_i_* associated with unit *i* of graph *G* completely bounds the feature **v**. The mapping is expressed as

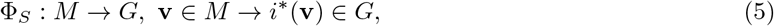

where the inequality

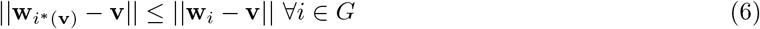

establishes the mapped vertex. The mapping Φ_*S*_ is topology preserving if adjacent features **v** ∈ *M* correspond to adjacent vertices of *G*, and therefore coincide with adjacent associated weights and resulting Voronoi polyhedra (Figure 4c). To satisfy this requirement, algorithm 1 is amended to include an additional step to adjust weights 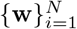 according to the neural gas algorithm [52]. The latter introduces an age *t_ij_* for each connection, and is used to remove elements *C_ij_* corresponding to weights, receptive fields, that are no longer adjacent following evolution. The final formulation of the TRN algorithm is:

(i) initialize each weight **w***_i_* for *i* = 1, ..., N, and set all connections *C_ij_* to zero;
(ii) present a pattern **v** ∈ *M*, where each **v** is drawn with equal probability;
(iii) for each *i* determine the number *n_i_* of units *j* where

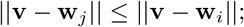
(iv) evolve **w***_i_* by the neural gas algorithm [52]

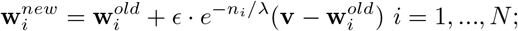
(v) If *C_ij_* = 0, set *C_ij_* > 0 (connect *i* and *j*) and set *t_ij_* = 0; else, leave *C_ij_* unchanged and set *t_ij_* = 0;
(vi) increment the age of all other connections made to unit *i*;
(vii) remove connections made to unit *i* that exceed a predefined age threshold; repeat at (ii).

**Figure 4:**
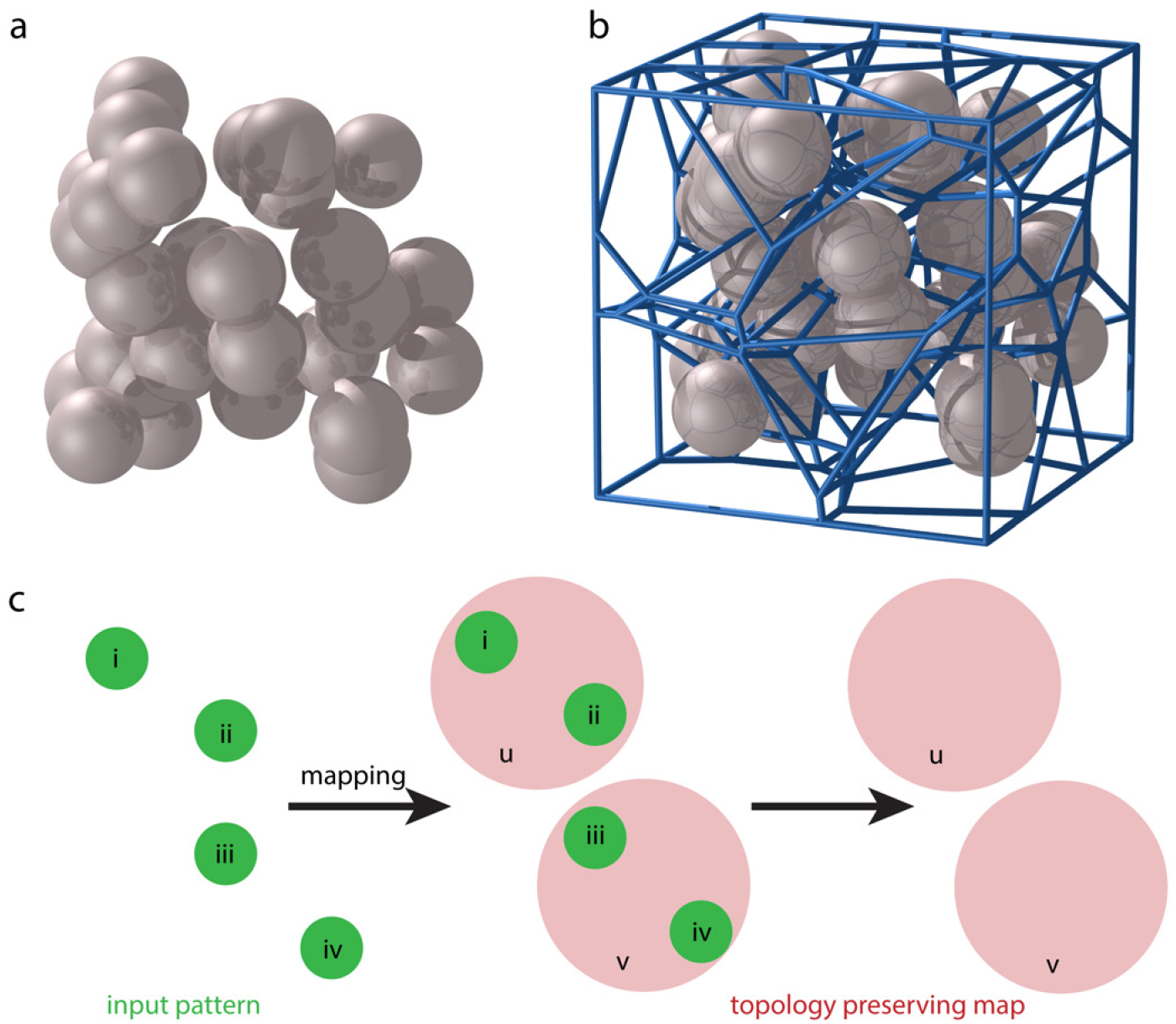
Topology preserving maps via 3-D Voronoi tessellation. **a** A 30-point cloud in 3-D, generated randomly in a cubic domain. **b** Resulting 3-D Voronoi tessellation of the point cloud in panel a. Each Voronoi cell partitions the spatial domain into regions that are closer to a given point than any other. Voronoi tessellation was performed with the *voro++* command-line tool [48] and rendering performed with the Persistence of Vision raytracer [49]. **c** Visual depiction and definition of a topology preserving map. The input pattern (green) consists of four points, enumerated i-iv. The coarse mapping groups two input points into a single coarse point (red), named u and v. The resulting mapping is topology preserving if adjacent features in the input pattern are adjacent in the output map. In this case, coarse point u maps to input points i and ii; coarse point v maps to points iii and iv; u and v bound adjacent groups of the input pattern and are adjacent in the output map.

Hyper-parameters *ϵ* and *λ* in step (iv) above are explained, as well as guidance for setting their values, in ref. [42].

In summary, the TRN computes a Delaunay triangulation from a set of weights that represent the locality of graph vertices, neural units, relative to features in the input space. For SBCG model building, each desired CG bead is treated as a neural unit, and its initial weight is a Cartesian coordinate within the embedding domain. Input patterns, i.e., atomic coordinates, are drawn sequentially from the reference molecule in a step-wise fashion. At each step, the weight associated with each neural unit is adapted until it is closer to its respective input pattern (atomic domain within the reference molecule) than any other. Movie M1 demonstrates the complete adaptation process using HIV-1 CA. To the best of our knowledge, the only widely-available implementation of this method is accessible via the CGBuilder plugin in the VMD molecular graphics software [53]. The latter is utilized in the present work. Post-optimization steps such as the assignment of atomic properties, as well as our approach to enable modeling in the sub-nanometer regime are discussed in the forthcoming sections.

#### Mapping of atomic properties to a SBCG model

Following optimization of the topology representing network, the properties of *N*_cell_ atoms within each Voronoi cell are mapped to CG beads. For a CG bead *j*, its mass *M_CG,j_* is computed according to

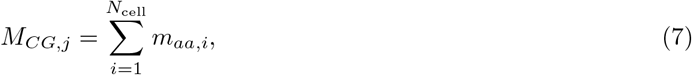

where *m_aa,i_* is the mass of atom *i* in the given Voronoi cell *j*.

Assignment of charge *Q_CG,j_* to CG bead *j* is analogous:

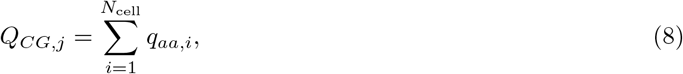

where *q_aa,i_* is the charge of atom *i* in the Voronoi cell *j*.

Non-bonded interaction terms, particularly the Lennard-Jones *ϵ*, well depth, parameter for each bead *j* are computed based on the solvent accessible surface area (SASA), *σ*, of the atoms within the Voronoi cell [44]. That is

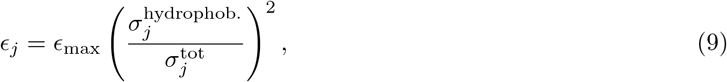

where 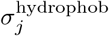 and 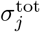 are the hydrophobic SASA and total SASA of the atomic domain in the Voronoi cell, respectively, and where *ϵ*_max_ is the maximum well-depth specified by the user. This formulation improves a previous formulation of the method where all beads are defined with a fixed *ϵ* value [43].

Finally, the last property assigned to CG beads following tessellation is the bead’s radius, which is important in properly representing the shape of the atomistic input. This is accomplished by computing the radius of gyration, *r*_gyr_, of the atomic domain with a given Voronoi cell.

Based on the above formulation, particularly equations 7 and 8, it is clear that such a CG reduction technique suffers loss-of-information in low-granularity use-cases. Information of the atomistic charge or hydrophobicity profiles, for instance, are critical for multimeric biological assemblies. In previous SBCG studies, assembly stability is maintained through specific, parameterized intermonomer interactions [45], ostensibly in the absence of detailed electrostatics and hydrophobicity information which are lost in the ∼150 atoms/bead mapping.

Utilizing the topology representing network aforementioned, available in VMD [53], requires the user to specify the granularity of the model, *N*_beads_, as a free parameter. On first inspection, it might seem trivial to increase the model granularity by simply increasing the *N*_beads_ parameter, and therefore prevent loss-of-information as outlined above. In practice, however, the available implementation fails to converge for any level of granularity finer than ∼40-50 atoms/bead (Figures S1 and S2). We addressed this issue with modifications to CGBuilder’s implementation of the TRN, which are detailed below.

#### Improving convergence of the topology representing network

Failed convergence of the topology representing network implemented in CGBuilder is caused by unintended *reflexive connections*, latent from how neuronal states are initialized prior to optimization. In learning theory, two neurons *a* and *b* are mediated by a reflexive connection if *a* ≡ *b*. That is, if *a* and *b* are indistinguishable to the optimization procedure, meaning their stimuli and associated scoring are identical, then their relationship is reflexive.

For the topology representing network implementation herein, we found that incident reflexive connections led to undefined behavior resulting in failed convergence. Specifically, in determining a graph representing a Delaunay triangulation via a *winner take all* selection rule, network behavior in the presence of a tie is undefined. If **w**_*i*_ = **w**_*j*_, the inequality

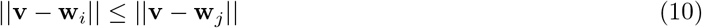

in step (iii) of algorithm 2 will evaluate identically for both *i* and *j*, leading to identical adaptation of **w**_*i*_ and **w**_*j*_ in the following step. Most critically, steps (v)-(vii) of algorithm 2 will determine *C_ij_* to be the strongest synapse, refresh its associated age *t_ij_* to zero, age every other connection made to *i*, then remove all other connections to *i* from **C** that exceed the age threshold.

Initialization of the network involves the instancing of one neuron per each of *N*_beads_. The input patterns are drawn from the atomistic structure [42], and the CGBuilder implementation pseudo-randomly selects Cartesian coordinates from the input to initialize the weights 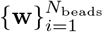 of neurons. During optimization, the weights are iteratively updated toward unique domains of input atoms (Movie M1) to which they are more proximal than any other (algorithm 2). For two neurons initialized with identical states, optimization forces them to identical final states. Only one bead will be assigned the properties of the atomic domain solved by the optimization and the remaining bead in the same cell will raise an error that is returned to the user, warning that there are no atoms to assign to the bead.

Our approach to enable higher granularity modeling enforces exclusivity among the initial states of neurons. During initialization, we maintain a record of which atoms among the input have already been utilized as an initial state. During pseudo-random selection of initial states, the record is conferred to assert that a given state has not yet been utilized, and if it has, we pseudo-randomly select another state. By enforcing an exclusivity condition while initializing the network, the optimization can be successfully applied to high-granularity use cases. In the following section, we will put forth a granularity selection criterion based on reciprocal space correlation between charge densities of the atomistic structure and resulting SBCG model, measured with Fourier Shell Correlation (FSC).

### Granularity selection via charge density Fourier shell correlation

Following our improvements to aid convergence of the topology representing network, we aimed to utilize a quantitative metric to motivate and establish a basis for selecting model granularity. To this end, we employ Fourier Shell Correlation (FSC) [54] between two charge density grids, one derived from the atomistic reference structure and the other resulting from SBCG mapping.

#### Computing charge densities

Charge densities are computed according to the charges on the molecular models, both atomistic and SBCG. For the present study, we employ the VolMap plugin in VMD [53]. First, the structures are cast to a 3-D voxel grid, with grid spacing of 0.5 Å. Each atom in the structure is modeled as a normalized Gaussian distribution, with distribution widths equal to the van der Waals radii of the atoms or beads. The Gaussians in the grid are then additively distributed. The resultant grids store charge density in 3-D space, which are amenable to FSC analysis. We employ the latter to gauge the fitness of our SBCG models to the atomistic reference from which it was derived.

#### Fourier Shell Correlation

Fourier Shell Correlation (FSC) is a commonly-employed method of measuring model-to-map fitness, map-to-map fitness and other correlation quantities in electron microscopy modeling [55, 56, 54]. The charge density grid represents a discretized real-space array *f*(**n**) where the domain **n** = (*n*_1_, *n*_2_, *n*_3_) corresponds to the Cartesian axes, and where each voxel in the grid stores a charge value.

In order to measure correlation between two charge density grids, their structure factors *F*(*r*) are first computed from the three-dimensional discrete Fourier transform (DFT) [57, 58]. For the spatial domain **n** = (*n*_1_, *n*_2_, *n*_3_) of extent **N** = (*N*_1_, *N*_2_, *N*_3_), the DFT convolves *f* (**n**) into the reciprocal spatial frequency domain *r* (Å^−1^) as

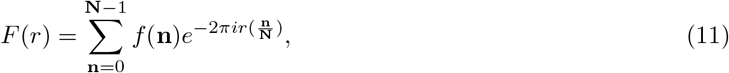

where 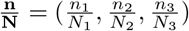 and where 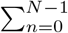 is the nested summation 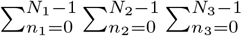.

Following convolution, the two charge density grids, denoted as *F*_1_(*r*) and *F*_2_(*r*), are subjected to FSC analysis. FSC measures a normalized cross correlation histogram, denoted here as *ζ*, across bins of increasing spatial frequency as

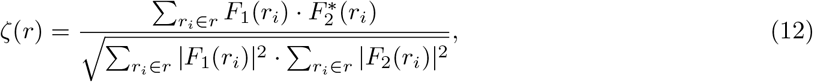

where 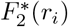 is the complex conjugate of *F*_2_(*r_i_*). Following calculation of *ζ*, we evaluate the histogram at specific correlation values as *ζ_n_*, where *n* is a number ∼[0, 1], to derive a model resolution. A value of *n* = 0 indicates that the structure factors are entirely uncorrelated, whereas *n* = 1 indicates perfect correlation between structure factors. Typically, the latter values of *n* are the so-called gold (0.143) and half (0.500) metrics. The assertion of resolution based on FSC will be elaborated in the following section.

#### Model selection based on charge density correlation

To assess the benefits of increased granularity with respect to accurately representing an atomistic charge density, we utilized our HIV-1 CA (Figure 5a,b), actin (Figure 5c,d) and cofilin-2 (Figure 5e,f) structures and computed a sets of SBCG models ranging from low to high granularity. The atomistic reference and each of the SBCG models were subjected to charge density calculation as outlined above, with special care taken to ensure that the van der Waals radii of the SBCG models were properly asserted before casting charges to the 3-D voxel grid (Figure 2c), since charge density depends on vdW radius (see *Computing charge densities*).

**Figure 5:**
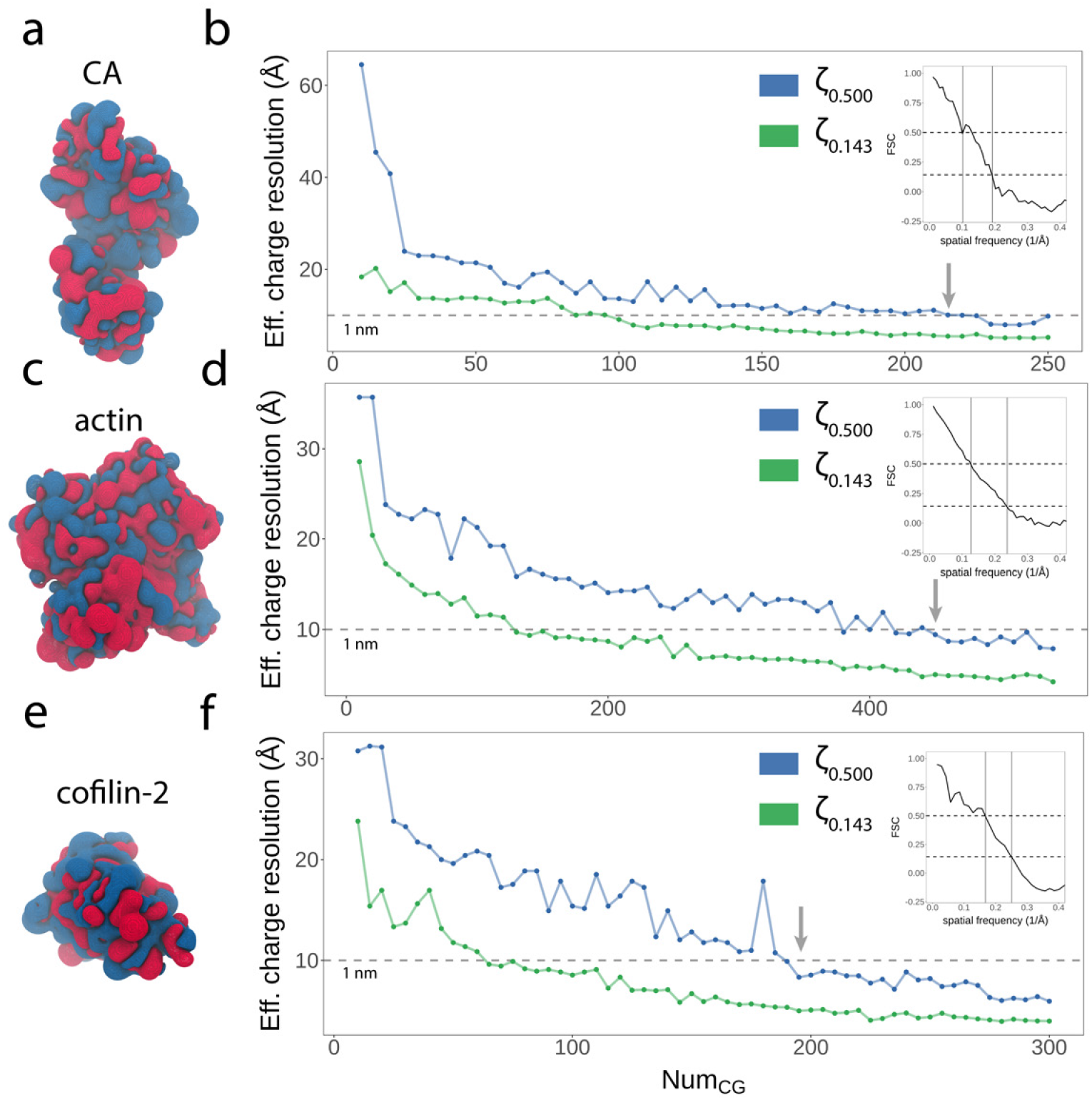
FSC analysis of CG granularity vs. effective charge density resolution. **a** Charge density of the all-atom reference structure of HIV-1 CA. Regions of positive and negative charge density are colored blue and red, respectively. **b** Effective charge density resolutions via FSC for models Num_CG_ ∈ [10, 250], plotted with two metrics: *ζ*_0.143_ and *ζ*_0.500_, green and blue, respectively. The dotted gray line represents a resolution of 1 nm, and the gray arrow indicates the first sub-nanometer model in the series. The inset plot shows the FSC vs. spatial frequency trace for HIV-1 CA with Num_CG_ = 250. **c** Charge density of the all-atom reference structure of actin and **d** corresponding effective charge density analysis for actin models with Num_CG_ ∈ [10, 540]. The inset plot corresponds to the FSC vs. spatial frequency trace for actin with Num_CG_ = 540. **c** Charge density of the all-atom reference structure of cofilin-2 and **d** corresponding effective charge density analysis for cofilin-2 models with Num_CG_ ∈ [10, 300]. The inset plot corresponds to the FSC vs. spatial frequency trace for cofilin-2 with Num_CG_ = 540.

To interpret the analysis, we employ the *ζ*_0.143_ and *ζ*_0.500_ metrics (Figure 5b, d and f), commonly-used to estimate resolution of particle reconstructions from electron microscopy. Metrics to determine reconstruction resolution are a subject of significant study and debate [59, 60]. In general, the FSC analysis considers amplitudes in structure factors at increasing radii of spatial frequency, or inverse resolution (Å^−1^) [56, 61]. The point along the spatial frequency axis at which correlation of two structure factors diminishes steeply is used for determination [62]. The two metrics, *ζ*_0.143_ and *ζ*_0.500_, have each been argued as effective methods of determining resolution, and additional analyses such as the ResLog plot have been put forth to ensure accurate particle alignment, free of aberrant correlation [60]. In our case, we are assessing the correlation among two charge density grids, where charges are interpolated from structures with differing granularity.

Our motivation for utilizing FSC is to assert an optimal granularity for representing the reference charge density with < 10 Å correlation, while adding as few degrees of freedom as possible and thus limiting computational expense of subsequent simulations. Trivially, a CG model with one representative bead per atom, and thus a one-to-one mapping of charge to each bead, would be perfectly correlated. Our results indicate that SBCG models for HIV-1 CA fall below 10 Å resolution in excess of 210 beads, employing the more stringent *ζ*_0.500_ metric. For actin, the first sub-nanometer model in the series was found to consist of 450 beads, and for cofilin-2, 195 beads. Figures S7, S8 and S9 show additional details of this analysis for CA, actin and cofilin-2, respectively, with examples of SBCG charge densities and additional FSC vs. spatial frequency traces.

Based on our analysis and subsequent determination of correlation of < 10 Å, we created a model of HIV-1 CA containing 221 beads, representing one bead per protein residue for the CA sequence utilized. For actin and cofilin, we chose 500 and 270 bead models, respectively, representing an approximately equal ratio of atoms to beads for each structure. While we construct and optimize SBCG models separately, the latter choice was made in anticipation of adjoining the models to constitute the heteromultimeric assembly. We then proceeded to the critical step of parameterizing the bond and angle terms governing the model, which are essential for accurately reproducing dynamics.

### Parameterizing sub-nanometer SBCG models

Parameterization of the SBCG bond and angle terms is accomplished with Boltzmann inversion, which is a technique commonly utilized [63, 43, 44, 45, 46, 47, 64]. Boltzmann inversion is employed to derive force constants based on mean square displacement (MSD, equation 14) of bonds and angles during all-atom simulation. For parameterizing HIV-1 CA, we employed an all-atom simulation of an HIV-1 CA trimer of dimers (Figure S4a), the latter constructed from six CA monomers. While we parametrize only a single SBCG CA monomer, the benefit of utilizing an assembly construct for inversion is threefold. First, the aggregate sampling of the atomistic trajectory totals nearly half a microsecond, 480 ns; second, the corresponding SBCG trimer of dimers (Figure S4b), simulated throughout iterative refinement, provides ample opportunity for cross-validation throughout the process; and third, to preserve the dynamical behavior of the CA monomers (Figure S4c) in their assembly environment given the state-dependence of Boltzmann inversion. For actin and colifin-2, we utilized a similar approach, performing all-atom simulation of a single globular actin bound to one human cofilin-2 protein [37].

In the following subsections, the formulation of Boltzmann inversion, the iterative refinement protocol (Figure 6a,b) and the necessary considerations for optimization of sub-nanometer structures, particularly removal of overlapping degrees of freedom (Figure 6c), are discussed.

**Figure 6:**
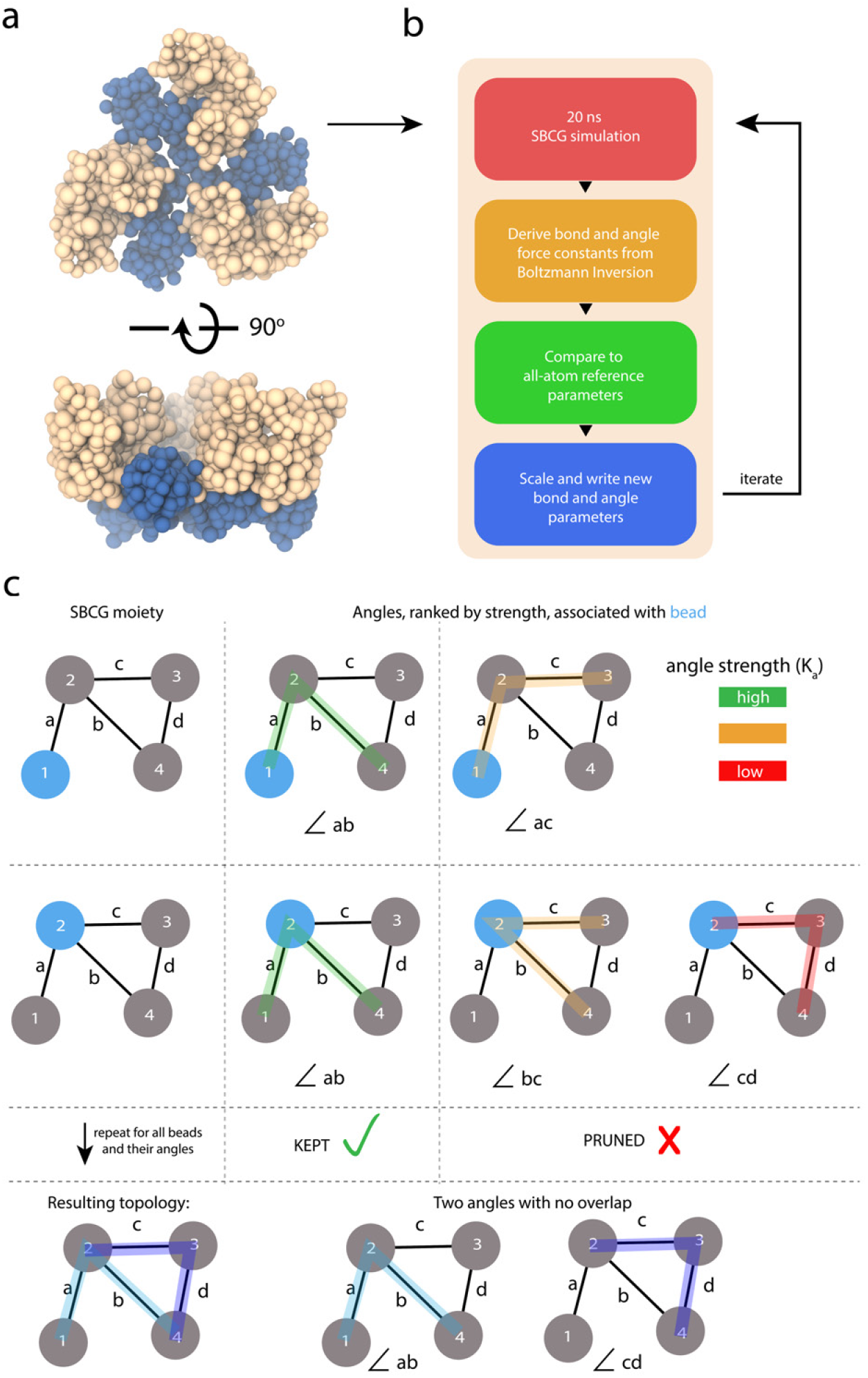
Graphical overview of our SBCG model refinement protocol. **a** SBCG HIV-1 CA trimer of dimers, utilized for successive 20 ns simulations during iterative refinement. The N-terminal domain is colored tan, and the C-terminal domain is colored blue. **b** The iterative parameter refinement procedure via Boltzmann inversion. For one iteration, 20 ns of equilibrium sampling is collected at 298 K. Next, bond and angle force constants are derived via Boltzmann inversion (equation 13). Parameters derived from SBCG are then compared to the all-atom reference parameters and are scaled by their error (equation 15). Finally, new bond and angle parameters are written and employed for the succeeding refinement iteration. **c** Graphical example of the pruning procedure employed in refining our model. This example shows a SBCG structure of four beads enumerated 1 through 4 and four bonds a through d. Initially, angles are determined exhaustively based on the bonded connectivity. For each bead, we rank its associated angle parameters by their constants, *K_a_*, and keep only the strongest parameter. This example demonstrates that our algorithm permits two beads to share the same force constant, if it is deemed the strongest for each bead.

#### Boltzmann inversion from atomistic simulation

Boltzmann inversion derives force constants for bonds and angles, *K_b_* and *K_a_*, respectively, according to

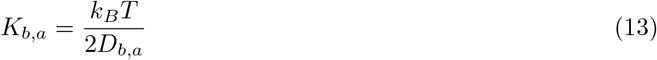

where

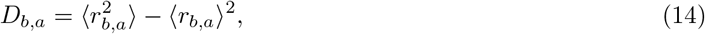

and where *r_b_* and *r_a_* are the measured bond and angle values. Units of *K_b_* and *K_a_* are *kcal mol* · Å^−2^ and *kcal mol · rad*.^−2^, respectively. *k_B_* is the Boltzmann constant and *T* is the absolute temperature, in units of Kelvin.

After initial derivation of bond and angle force constants from the all-atom simulation, we performed a SBCG simulation with the resulting parameters, and utilized Boltzmann inversion targeting the SBCG trajectory as validation; we observed a terrible fit (Figure 7 and S10a). This behavior of the Boltzmann inversion method has been reported elsewhere [45, 47] and is a known short-coming of this approach. The problem is in the assumption that each bond and angle are independent. In reality, bonds and angles are highly coupled throughout the structure and this is especially true in the sub-nanometer SBCG regime. To remedy this, we employ an iterative refinement protocol, based on previous work [47].

**Figure 7:**
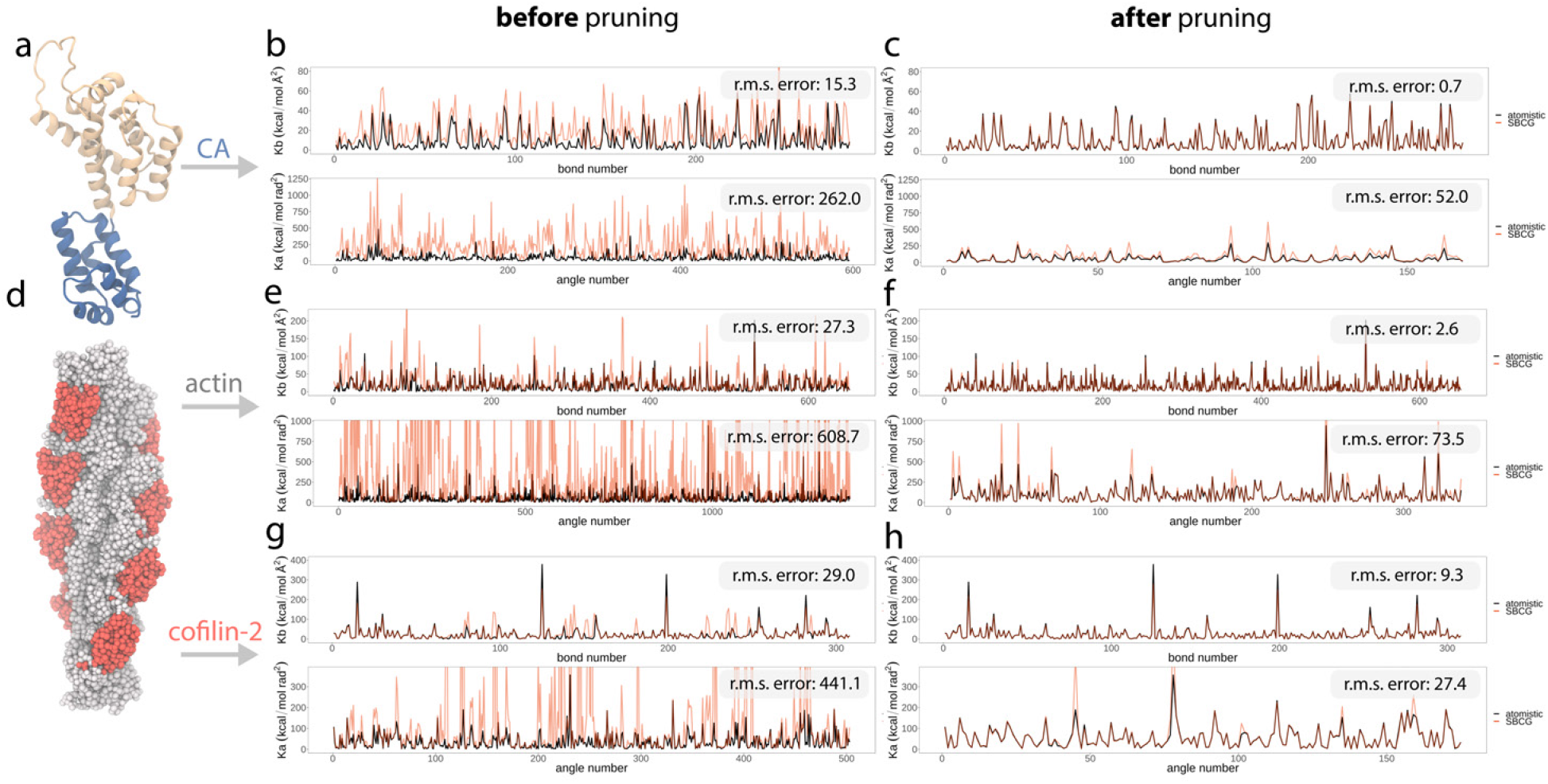
SBCG parameter optimization results for HIV-1 CA, actin and cofilin-2. **a** Monomeric HIV-1 CA all-atom structure, shown in cartoon representation. **b** and c Corresponding bond and angle parameter fits following from iterative Boltzmann inversion. The black traces show the atomistic bond and angle parameter trace as computed via Boltzmann inversion from the all-atom reference trajectory, and orange the SBCG parameter trace via Boltzmann inversion from the simulation corresponding to the final refinement iteration **b** before pruning and **c** after pruning. **d** Atomistic surface representation of cofilin-2 bound to one turn of actin. **e**,f Actin bond and angle parameter fits following from iterative Boltzmann inversion **e** before pruning and **f** after pruning. **g**,**h** Cofilin-2 bond and angle parameter fits following from iterative Boltzmann inversion **g** before pruning and **h** after pruning. The root-mean-square error between SBCG parameters and their respective all-atom reference parameters are annotated within each plot.

#### Iterative refinement

From refinement iteration *i*, the parameters for the next iteration *i* + 1 are computed according to

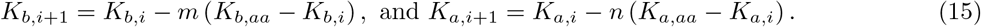

*K_b,aa_* and *K_a,aa_* are the bond and angle force constants derived from the all-atom reference trajectory. The constants *K_b,i_* and *K_a,i_* are derived from Boltzmann inversion of 20 ns SBCG simulations. Variables *m* and *n* are scaling constants, and are treated as hyperparameters [47].

Prior to deploying the above protocol, we performed a parameter sweep to identify optimal *m* and *n* scaling parameters. The sweep covered *m* and *n* ∈ [0.1, 0.9] with a stride of 0.1 for each constant, resulting in 81 separate SBCG simulations 20 ns in length. Inversion was then applied to these trajectories to yield parameters, and the improvement from the previous parameter set was measured via root-mean-square error (RMSE) (Figure S5).

With optimal *m* and *n* scaling constants identified, we performed many iterations of refinement. After, it became clear that the parameters had improved, but converged to an unphysical state with poor fit (Movie M2). For all three of our structures, angle parameters were particularly problematic. We determined that the problem is caused by high connectivity, and therefore redundant degrees of freedom (Figure 7b, e and g, Figure S10b,c).

#### Pruning redundant degrees of freedom

Figure S10 shows the analysis utilized to identify the cause of unphysical convergence. For each angle parameter, comprised of three CG beads, we analyzed the connectivity associated with the beads. Angle parameters with the poorest fit were found to involve beads with high connectivity. Conversely, we found that angle parameters involving CG beads with relatively few bonded terms were well-fit. Regions exemplary of the latter are shaded with red and green, respectively, in Figure S10c.

Further, we analyzed violations, i.e., deviations of the SBCG angle vs. its all-atom reference value, and found that behavior of a given CG bead, and therefore its bond and angle parameters, is dominated by its strongest connections; weak parameters are overpowered by stronger, coupled parameters and thus a violation is manifest. The latter, coupled with regions of the topology containing many overlapping, redundant degrees of freedom, led to an untenable optimization problem. To remedy this, we collected for each CG bead the angle parameters with which it is associated. For each bead, only the strongest angle parameter (strongest meaning the highest force constant based on all-atom reference simulations) was retained. We refer to this process as *pruning* (Figure 6c).

#### Converged fit

Following pruning of redundant angle parameters, our optimization immediately converged to a better fit for all three of our structures (Figure 7c, f and h). While not scale-invariant, we employ root-mean-square error (RMSE) as a progress indicator of the fitting. The plots in Figure 7 are annotated with the bond and angle RMSE for each of our three structures, before and after pruning, quantifying how crucial removing redundant degrees of freedom is. Prior to any sub-nanometer SBCG parameter optimization endeavor, pruning should be performed because optimization, based on the present formulation where bonds and angles are treated independently, is otherwise untenable in high-granularity cases, as we have demonstrated with three unique structures. Our pruning algorithm (Figure 6) is available for easy-use within the CGBuilder plugin, distributed with VMD [53].

### Macromolecular assembly simulations

With the models parametrized, we proceed to constructing our macromolecular assemblies. Generally, applying a monomeric model to a multimer involves transferring the SBCG mapping of a single monomer to each subunit in the assembly. A critically important detail at this stage is that the subunit subjected to the initial CG reduction is identical in sequence and structure to those comprising the assembly.

In the following sections, the multimeric assembly mapping procedure will be discussed. Further, with the goal of including inositol hexakisphosphate in our SBCG conical capsid model, we will elaborate guidelines for including CG ions and small molecules, and highlight the importance of performing counter ionization via the model’s Coulombic potential, the latter step which is critical in the sub-nanometer model regime due to increased charge fidelity (Figure 5). Finally, we will discuss simulation configuration; and importantly, determination of the integration time step via calculation of model bond frequencies.

#### Extension of SBCG model to heteromultimeric assemblies

In SBCG modeling, the CG mapping refers to the atomic domain assigned to each bead following spatial tessellation according to the topology representing neural network (detailed in section *Shape based coarse graining*). Recall that each Voronoi cell emergent from network optimization bounds a domain of atoms, and the CG bead located at the center-of-mass of this domain is assigned the properties of constituent atoms. The multimeric mapping, or map transfer, protocol contained in the CGBuilder implementation, utilizes the information of the CG mapping to locate each domain, Voronoi cell, in equivalent atomistic subunits comprising the assembly. Topology and parameters, e.g., bonds, angles, mass, charge, derived in previous steps are copied to the new CG subunits.

For the map transfer operation to be successful, each target atomistic subunit must be identical in sequence and structure to the original structure employed for CG reduction. The necessity for equivalence manifests from the identification of atomic domains mapped to each bead. If these domains are in different spatial locations, then bonds, angles joining them will be violated when topology and parameters are copied to the new subunit. Additionally, if differences in sequence are present, then the map transfer as implemented may fail completely, or place beads in positions not intended by the user.

We recommend performing separate CG mappings and parameterizations for unique structures, if the assembly is heterogeneous. For homomultimeric assemblies, taking care to construct a target atomistic assembly from identical subunits will bypass this problem entirely. Proprietary, or in-house, alignment protocols may further be employed as a solution to mapping to similar, but not equivalent, structures.

#### Coarse grained ions and small-molecules

CG flavors and force fields have differing ways of treating ionic, or otherwise charged, species. In the present study, we chose an approach following the work of Arkhipov, et al. [46], which describes anionic and cationic species, chloride and sodium, as groups of ions which carry either a −1 or +1 charge, respectively. Arkhipov modeled these as groups of nine ions. Given the increased granularity of our models, we chose a similar approach but utilized smaller groupings of ions, five in total for both positive and negatively charged ions. The latter choice was made according to the largest bead, by mass, in our SBCG protein topology. In our testing, inclusion of ionic species with vastly different mass than that of protein beads caused numerical instability when attempting to utilize large integration time steps.

Our SBCG conical capsid model includes an additional, small molecule species: inositol hexakisphosphate, or IP6 (Figure S3a). IP6 is a highly charged molecule, at −12 *e*, and a known assembly co-factor for HIV-1 capsids [39, 40]. In our model system, IP6 is treated as a single bead of radius 5 Å (Figure S3b). CG IP6 was assigned a charge of −12 *e*, and 253 were placed corresponding to the 253 capsomers comprising the conical capsid (Figure S3c). In atomistic HIV-1 CA hexamers and pentamers, IP6 resides approximately perpendicular to the Arginine 18 ring situated at the central pore [65]. We utilized this information to place CG IP6 beads in our model (Figure S3c).

#### Counter ionization via 3-D Coulombic potential

As we have pointed out, sub-nanometer CG models have high charge fidelity, and it is therefore necessary to balance the charges of the initial model with counter ions, similar to preparation of an atomistic model. To this end, we employ a Coulombic grid potential calculation available in VMD [53] named CIonize. In a discretized 3-D grid, Cionize computes a Coulombic potential iteratively after successive placements of ions. Interestingly, and perhaps serving as an additional validation of the detailed charge of our HIV-1 CA model, Coulombic potential calculations placed Sodium and Chloride in equivalent positions to where these ions are known to reside in atomistic resolution structures [41] (Figure S6).

With charges balanced, and other considerations addressed such as the inclusion of small molecules or cofactors, we now turn our attention to configuring CG molecular dynamics simulations.

### Simulation parameters: temperature control, time step selection, long-range electrostatics

In the present study, we employ the NAMD molecular dynamics engine [38] for all simulations, for both optimization of parameters (Figure 6 and 7) and production simulations of our multimeric assemblies, i.e., the HIV-1 capsid and cofilin-2-bound actin filaments (Figure 1, 2 and 3). As with configuring an atomistic simulation, the selection of configuration parameters is a critical step in ensuring the physical realism of the resulting MD ensemble. Here, we place particular emphasis on temperature control, integration time step, electrostatic evaluation and the cut-off scheme; which, in a sub-nanometer context, have additional importance compared to low-granularity CG models.

#### Integration time step

Among the most fundamental choices when configuring a molecular simulation is the value of the integration time step. In most circumstances, choosing an integration time step—and thus establishing the temporal resolution—is motivated by the scale of the system, atomic or otherwise. For instance, an atomistic simulation of a protein might employ a 1-2 femtosecond (fs) time step, small enough to capture vibrational modes of a bond to hydrogen (∼10 fs). In practice, bonds to hydrogen may be constrained and access to larger, or multi-timescale, integration steps becomes possible. This confers better computational performance, increasing sampling and broadening the temporal resolution of the ensemble such to capture collective, large-scale molecular motions.

In SBCG modeling, and particularly sub-nanometer SBCG modeling, the selection of the time step is determined based on two factors: the masses of the CG beads comprising the model, and the force constants, *K_b_*, employed in the bonded potential energy terms. Because SBCG does not follow a mapping scheme a priori, but rather computes a mapping through neural network optimization, time step selection depends on granularity, more specifically the resulting CG bead masses, and is motivated by evaluating vibrational frequencies in the model.

For a bond *i*, vibrational frequency *ν_i_* is computed according to

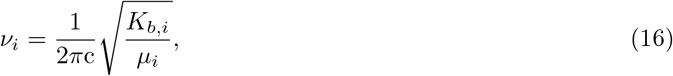

where *c* is the speed of light, *K_b,i_* is the bonded force constant and *μ_i_* is the reduced mass of the two beads involved in the bond

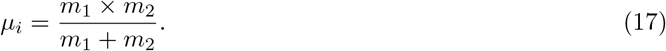

Following evaluation of vibrational frequency for all bonds comprising the CG topology, the time step *τ* is then taken from the set of all frequencies 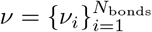 as

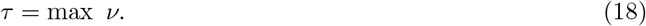

That is, we compute vibrational frequencies for the complete topology and choose a time step based on the fastest vibration, i.e., smallest oscillation period, present. Because the bonded force constants are optimized during iterative refinement, we recommend first evaluating equation 18 using the initial parameter set yielded by Boltzmann inversion (equation 13) of the atomistic trajectory, then re-evaluating following iterative optimization. Far-exceeding the fastest vibrational frequencies with the selected integration time step leads to numerical instability.

#### Temperature control

For all SBCG simulations, we sample constant temperature (NVT) ensembles with temperature control via Langevin dynamics. The latter controls temperature by coupling the particles in the system to a dissipative background force and a randomly fluctuating force. Specifically, for a particle with mass *m* and position **x**, subjected to dissipative force **f** (**x**) = −∇*U* (**x**), its motion is [66]

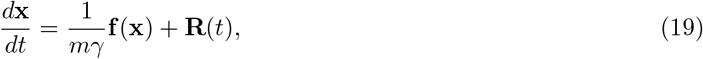

where **R** is a zero-mean, Gaussian random process [67] such that

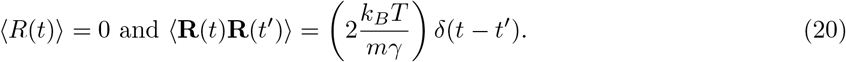

Importantly, the coefficient *γ*, in units of inverse time, is a user-specified parameter that controls the strength of thermal coupling; this is also referred to as a friction term. In NAMD’s stochastic formulation of Langevin dynamics [67, 38], the dissipative and fluctuating force terms in equation 19 are added to the Newtonian equations of motion to achieve thermal coupling and thus temperature control. Importantly, the choice of the Langevin *γ* term has special significance to the dynamical evolution of the molecular system [68].

Temperature control in the Langevin framework relies on several considerations, the most principal of which is the intended dynamical regime. In molecular dynamics, momentum is conserved and inertial effects of particles are significant. In Langevin dynamics, dampening of velocities, and thus momentum, through coupling to an external thermal reservoir – introducing a stochastic differential equation to Newton’s equations of motions [69] – allows temperature control. Increasing the *γ* coupling parameter, the system tends toward the overdamped limit [68], where inertial effects are diminished and Brownian dynamics begin to dominate. In the Brownian dynamics regime, momentum is not conserved [68]; particles comprising the system feel a random force and a drag force, or friction, relative to a constant background (eq. 19), and thus their motions become Brownian [68, 70].

In several CG modeling contexts, we note the reported use of *γ* coefficients in excess of 10 − 100 ps^−1^, whereas in atomistic molecular dynamics contexts, *γ* is typically held between 0.5 − 2.0 ps^−1^. It is worth noting that overdampening is one method of achieving numerical stability during simulation, granting access to larger integration timesteps. We caution the reader against indiscriminately increasing their friction coefficient to dampen velocities, unless they are aware of the dynamical consequences. For instance, performing self-assembly simulations is one exemplary justification for overdampening and accessing a larger integration timestep.

In our systems of the HIV-1 SBCG conical capsid as well the cofilin-2-bound actin filaments we employ a *γ* of 2.0 ps^−1^, primarily to model, implicitly, the viscosity of water. Throughout testing we observed that we could make our time step arbitrarily large by increasing *γ* indiscriminately. Achieving a large time step is desirable only from the vantage of computational performance. If increased sampling efficiency comes at the expense of the intended dynamical regime, or predictive capability, then we argue that this is not a worthwhile exchange. For SBCG molecular dynamics, a *γ* between 0.5 − 2.0 ps^−1^, in concert with an appropriate dielectric, will productively introduce some of the macroscopic effects of solvent, namely viscosity and charge screening.

#### Long range electrostatics via Particle Mesh Ewald (PME)

An additional, important consideration for the simulation of sub-nanometer SBCG models is the treatment of long-range electrostatics. One oft-utilized technique in molecular simulation is the Particle Mesh Ewald (PME) approach [71, 72]. In PME electrostatic evaluation, charges are interpolated on a discrete grid, or mesh, to compute the electrostatic potential. This method is parallelizable and has been described in detail, and the specific implementation employed in the NAMD molecular dynamics engine has similarly been described [67, 38]. We employ PME to treat long-range electrostatics in sub-nanometer SBCG simulations.

Utilizing PME in MD simulations confers detailed electrostatic treatment at the expense of performance. In our testing of the HIV-1 conical capsid, PME reduces the performance of our simulations by an approximate factor of four compared to truncated dynamics without any long-range electrostatic component (Figure 1d,e). The resolution, in Å, of the grid to which charges are cast is a free parameter. We have found that a grid resolution of 2 Å with a corresponding interpolation order of eight allows us to recover some of the lost performance, without sacrificing accuracy or numerical stability. Selection of grid resolution, and interpolation order, are use-case specific considerations. Further, the choice of electrostatic cutoff distances is an associated dependency in treating long-range electrostatics, which is discussed in the following section.

For the cofilin-2-bound actin filaments, we find that PME electrostatic evaluation leads to a more significant reduction in performance (Figure 2d,e) compared with truncated dynamics. The reason for this is related to the spatial decomposition of the filamentous systems, which have significantly large ratios of length to cross-sectional area. This point is notable, since SBCG modeling pushes molecular simulation to considerable size scales. We are motivated to address the latter in future work.

#### Non-bonded interaction cutoffs

Related to establishing parameters for the PME grid is the assertion of cutoff distances. In the NAMD molecular dynamics engine, the cutoff scheme is described with three parameters: a cutoff distance, beyond which the long-range potential is truncated; a switching distance (if *switching* is enabled in the configuration), which specifies the distance beyond which a splitting function is employed; and the pairlist distance, which determines the maximum considerable pair distance between any two particles.

Fundamentally, cutoff distances should be larger than the longest bond term in the CG topology; however, increasing electrostatic cutoff distance leads to larger computational expense, since more bead pairs in the pair list necessitate more evaluations. Utilizing the longest bonded distance in the topology as a lower bound, we employ an upper bound based on the interfacial distances in our biomolecular assembly. This approach is equivalent to an approach used to select cutoff distances in a previous SBCG study of capsids [45].

#### GPU accelerated coarse grained simulations

The sampling efficiency of SBCG simulations benefits significantly from GPU acceleration. Typical GPU accelerators have their own dedicated memory of 8 to 24 GB as of the time of this publication. In the GPU-accelerated computing paradigm, problems that fit neatly within the memory of the graphics processor are amenable to multiple-factor speed ups [73]. In contrast, problems that exceed dedicated accelerator memory lead to costly host-to-device copy operations and excessive communication overhead which place a hard limit on attainable sampling efficiency. The design strategy of MD engines such as NAMD2 [67] is to offload only a subset of computations to the GPU, namely evaluation of non-bonded electrostatics. While selective offloading is a flexible strategy that accommodates diverse systems on heterogeneous architectures, the biggest performance gains remain unrealized.

Recently, a fully GPU-resident MD engine NAMD3 [38] was developed, which offloads all computations to the GPU. Multimeric SBCG assemblies, such as the HIV-1 conical capsid presented here, represent ideal memory footprints for saturating and taking full advantage of GPU acceleration. Remarkably, with certain simulation configurations such as those employing truncated dynamics (see section *Electrostatics via Particle Mesh Ewald*), we are able to achieve sampling efficiency in excess of 1 microsecond per day using NAMD3 (Figure 1e) for the HIV-1 capsid, and greater than 3 microseconds per day for our 3-turn filament system (Figure 2e). Employing full electrostatic evaluation with PME for simulations of the HIV-1 capsid, we can still reach high sampling rates in excess of 300 nanoseconds per day (Figure 1d). The latter two performance metrics represent significant speedups over CPU, or heterogeneous CPU and GPU, computation. Furthermore, our benchmark analysis shows that the performance of multimeric SBCG assemblies scales across multiple GPUs. Table S1 shows benchmarks of the 3-turn cofilin-2 bound actin filament system utilizing NVIDIA’s DGX A100, employing varying numbers of cores per GPU utilized. Remarkably, utilizing eight A100 GPUs with eight CPUs per GPU, yielding 64 in total, we exceed four microseconds per day simulation performance.

## Conclusions

We have shown the utility of shape-based coarse grain (SBCG) modeling for efficient simulation of large biomolecular assemblies and have outlined the protocol for effective deployment. Further, we addressed the selection of model granularity via Fourier Shell Correlation analysis targeting atomistic and SBCG charge densities. Optimization of parameters, as well as removal of redundant degrees of freedom, was outlined and illustrated in detail such to reproduce atomistic behavior. We described numerous considerations for configuring and performing simulations of biomolecular assemblies using sub-nanometer SBCG, such as temperature control, computation of the integration timestep, and long-range electrostatics. Our code is freely-available as part of the CGBuilder plugin in VMD 1.9.4 [53], which is distributed with a corresponding tutorial and example files.

## Acknowledgements

The authors acknowledge funding from the US National Institutes of Health award U54AI170791 (to J.R.P.). This work used the Extreme Science and Engineering Discovery Environment, which is supported by the National Science Foundation (Grant ACI-1548562). This work used XSEDE Bridges and Stampede2 at the Pittsburgh Super Computing Center and Texas Advanced Computing Center, respectively, through allocation MCB170096.

## Supplementary Information

**Table S1:** Performance benchmarks of the 3-turn cofilin-2-bound actin filament system utilizing NVIDIA’s DGX A100, with 640 GB of total memory. On the left, performance metrics are obtained for SBCG MD simulations with PME *off*. On the right, performance metrics obtained for SBCG MD simulations with PME *on*. For each number of GPUs employed, the number of CPUs providing optimal performance is shown in bold.

| DGX A100 640GB |  |  |  |  |  |  |
| --- | --- | --- | --- | --- | --- | --- |
| No PME |  |  |  | PME |  |  |
| GPUs | CPUs per GPU | ns/day |  | GPUs | CPUs per GPU | ns/day |
| 1 | 8 | 1497.62 |  | 1 | 8 | 481.657 |
| **1** | **16** | **1564.61** |  | **1** | **16** | **496.476** |
| 1 | 32 | 1539.68 |  | 1 | 32 | 491.068 |
| 1 | 64 | 1474.44 |  | 1 | 64 | 485.423 |
| 1 | 128 | 1327.73 |  | 1 | 128 | 442.508 |
| 2 | 8 | 2305.98 |  | 2 | 8 | 548.472 |
| 2 | 16 | 2370.77 |  | 2 | 16 | 549.247 |
| **2** | **32** | **2433.6** |  | **2** | **32** | **552.919** |
| 2 | 64 | 2265.39 |  | 2 | 64 | 532.441 |
| 4 | 8 | 3111.83 |  | 4 | 8 | 595.852 |
| **4** | **16** | **3247.3** |  | **4** | **16** | **598.493** |
| 4 | 32 | 3225.44 |  | 4 | 32 | 597.041 |
| **8** | **8** | **4038.3** |  | 8 | 8 | 621.336 |
| 8 | 16 | 3711.83 |  | **8** | **16** | **621.695** |

**Figure S1:**
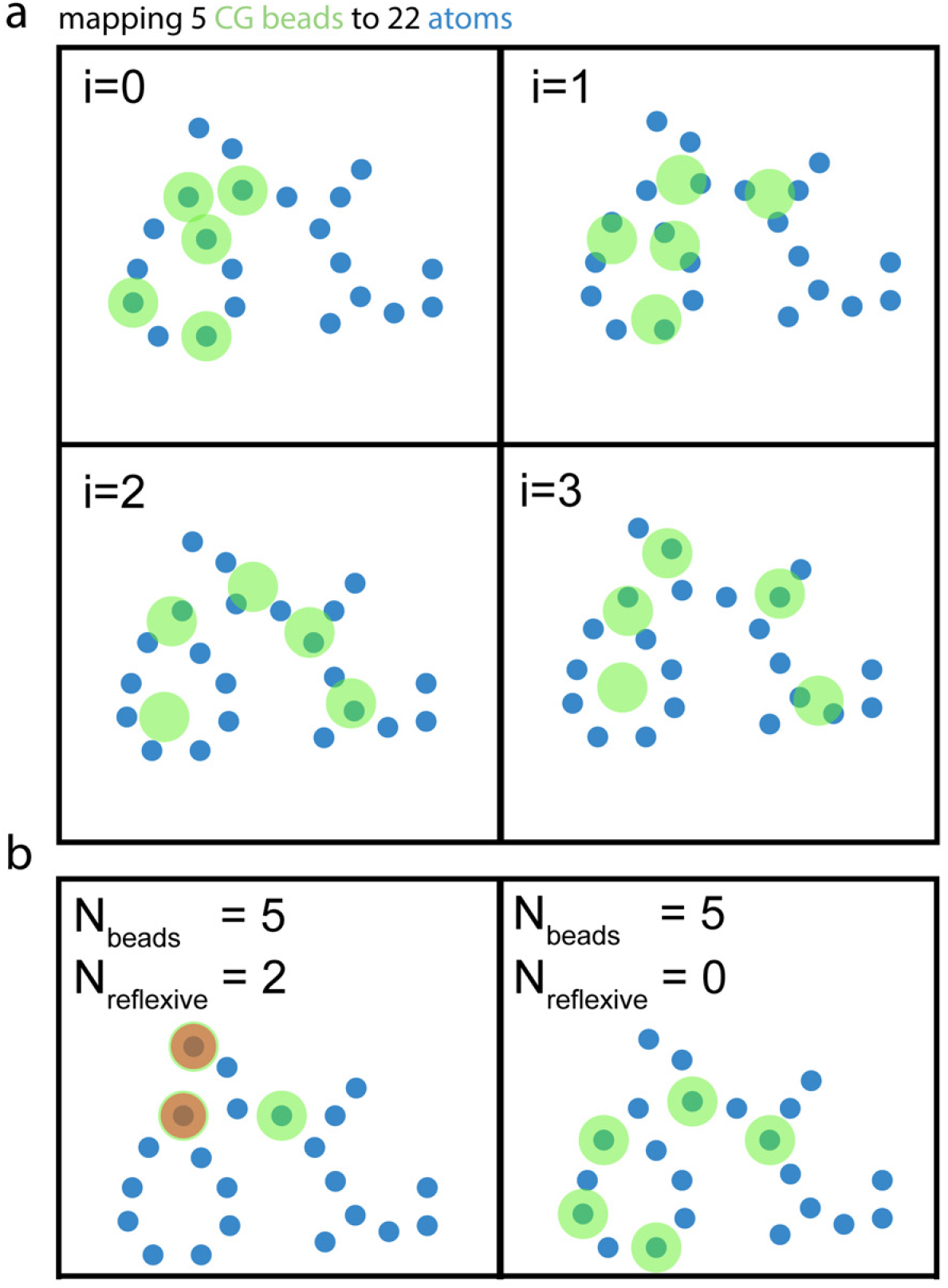
Graphical overview of the topology representing network with a 2-D cartoon example. **a** Exemplary 2-D graphic showing the mapping of five CG beads (green) to 22 atoms (blue), over four iterations, or *learning steps*, *i*. **b** Left, example case of unintended *reflexive connections* (red), where two CG beads are initialized to the same coordinate prior to network optimization. This case results in two assignment failures. Right, the same example but where each bead and its associated weight begin at a unique starting position.

**Figure S2:**
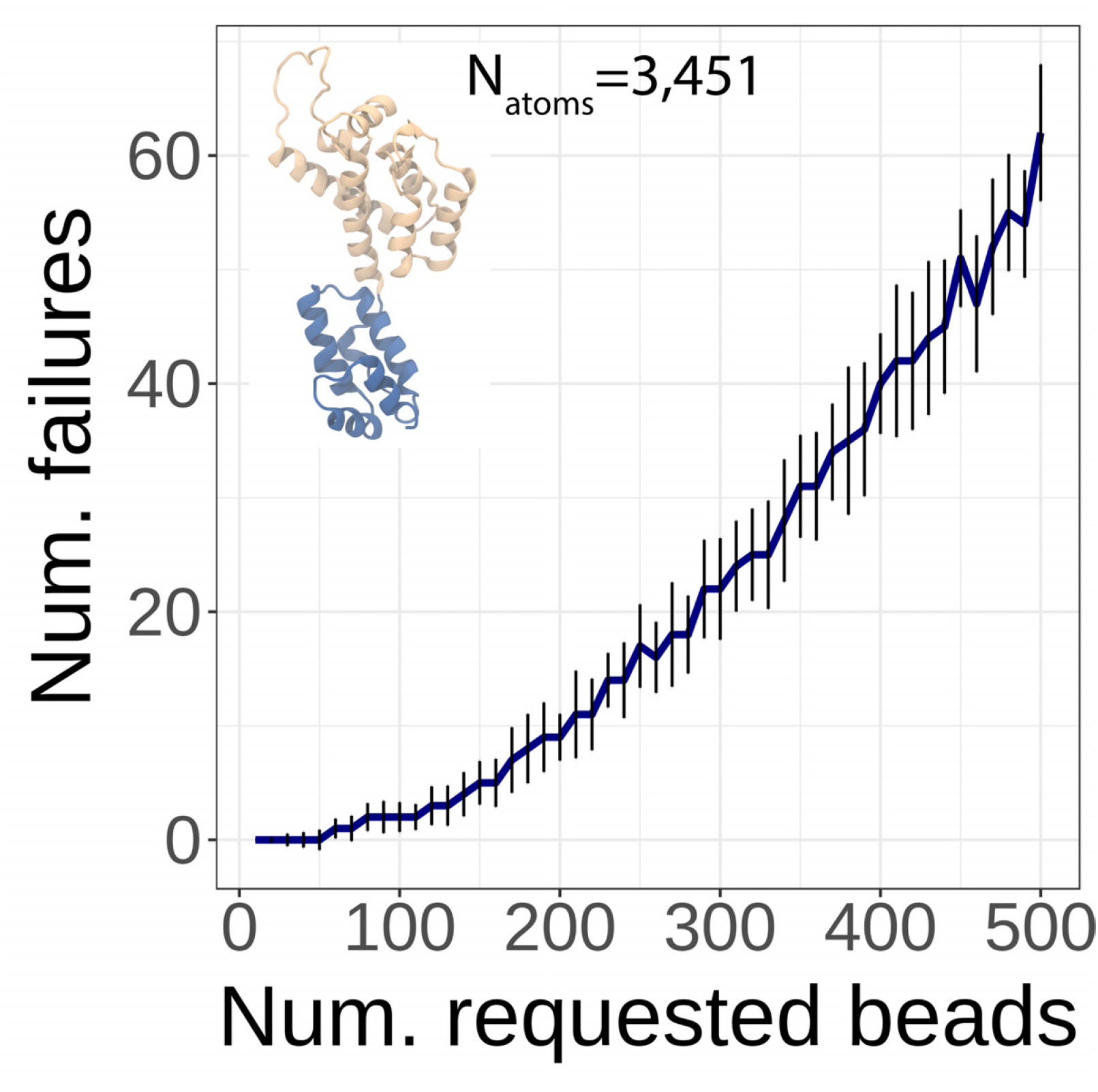
Assignment failures in the Topology Representing Network (TRN) resulting from reflexivity. Without an exclusivity condition among neuronal weights, runs of the TRN produce assignment failures as illustrated in Supplemental Figure S2. To perform this analysis, the relevant code was extracted from the original CGBuilder plugin and written into a simulator, which for a given input pattern (in this case HIV-1 CA, shown inset) can determine how many assignment failures will manifest. The analysis performed tests over *Num*_CG_ ∈ [10, 500] with a stride of 10 for an input of 3,379 atoms, and each value of requested beads was simulated 20 times. The error bars denote the standard deviation over these 20 tests, the latter deviation among the ensembles results from psuedo-random selection.

**Figure S3:**
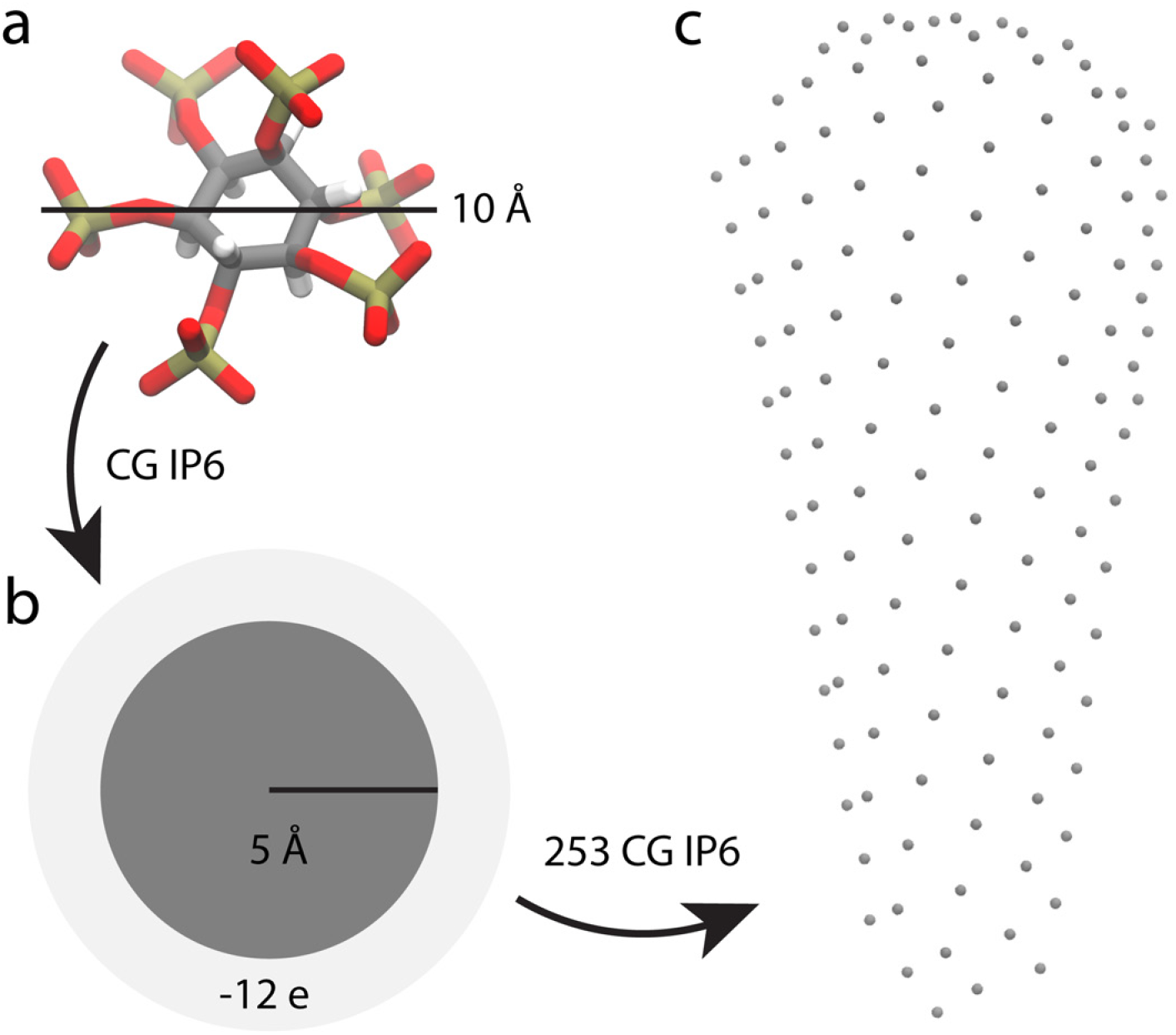
Overview of the CG description and placement of inositol hexakisphosphate (IP6) assembly cofactor. **a** Atomistic IP6 model, measured with a diameter of 10 Å. **b** Single-bead CG IP6, parametrized with a diameter of 5 Å and a charge of −12 *e*. **c** Final placement of CG IP6 beads in our conical capsid model. One CG IP6 is placed at the center pore of each capsomer, yielding 253 in total. The beads are placed proximal to beads mapped to side chains of Arginine 18.

**Figure S4:**
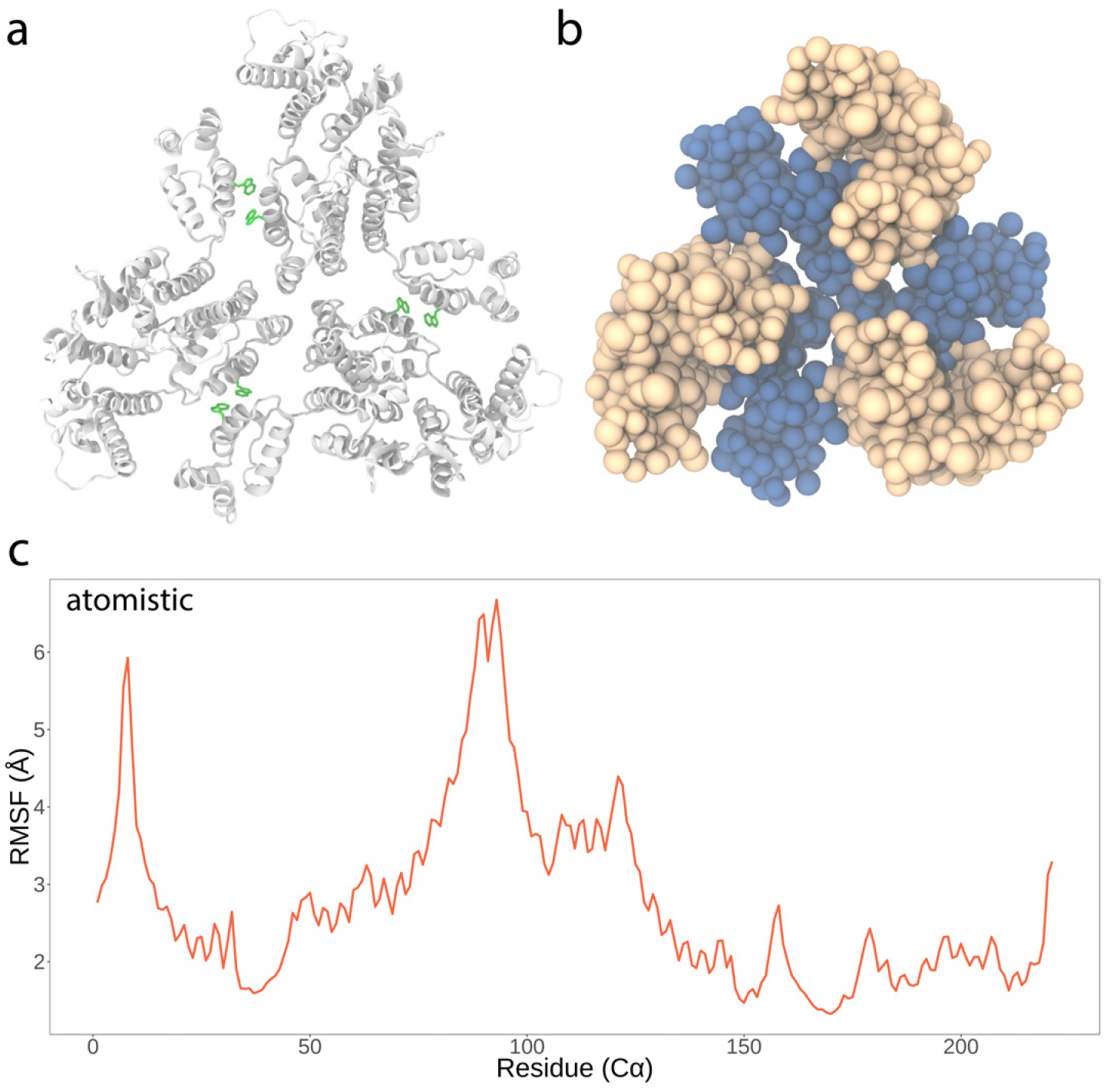
HIV-1 CA trimer of dimers construct utilized in parametrization. **a** Atomistic CA trimer of dimers, simulated at 298 K for 80 ns. Protein is shown in cartoon representation. Tryptophan 184, critical for intermolecular dimer interface stability, is shown in green licorice representation. **b** Resulting SBCG trimer of dimers, which was simulated throughout iterative refinement (Fig. 6 and 7). **c** Root-mean-square fluctuation (RMSF) of alpha Carbons of the atomistic construct in panel a.

**Figure S5:**
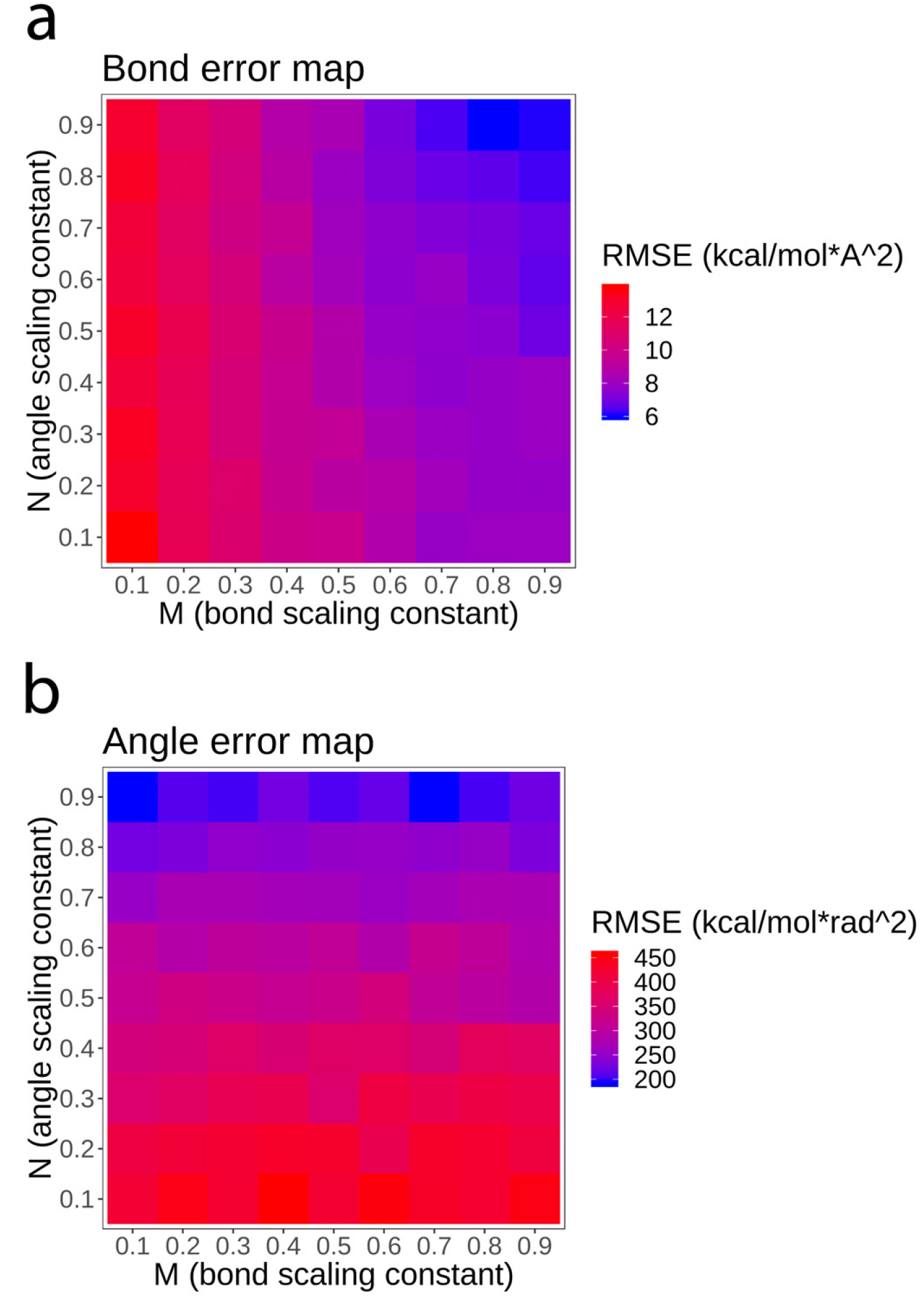
Parameter sweep for identifying optimal *m* (bonds, **a**) and *n* (angles, **b**) scaling constants. The heat map shows the root-mean-square error for 81 simulations – a 9×9 grid – computed for a single refinement iteration. Blue denotes lower error and red denotes higher error, with scale bars provided.

**Figure S6:**
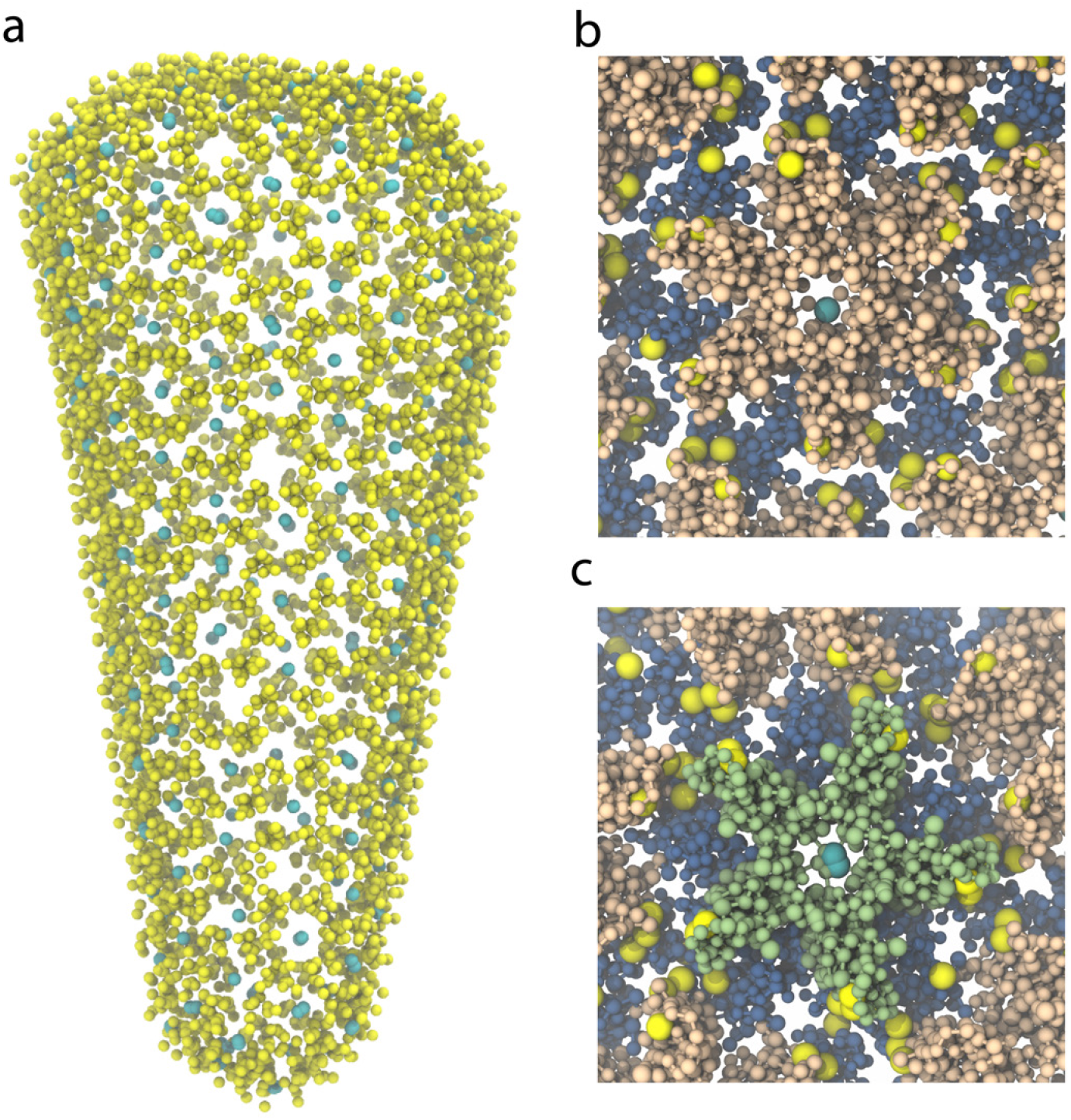
Counterion placement around the SBCG HIV-1 conical capsid. Radii of protein and ion beads are not to scale, but selected for visual clarity. **a** Counterions of mixed nature, carrying either a positive (yellow) or negative (cyan) charge, as placed according to iterative Coulombic grid potential calculations. **b** Close-up of a CA hexamer from the conical capsid, showing counterions. **c** Close-up of a CA pentamer from the conical capsid, showing counterions.

**Figure S7:**
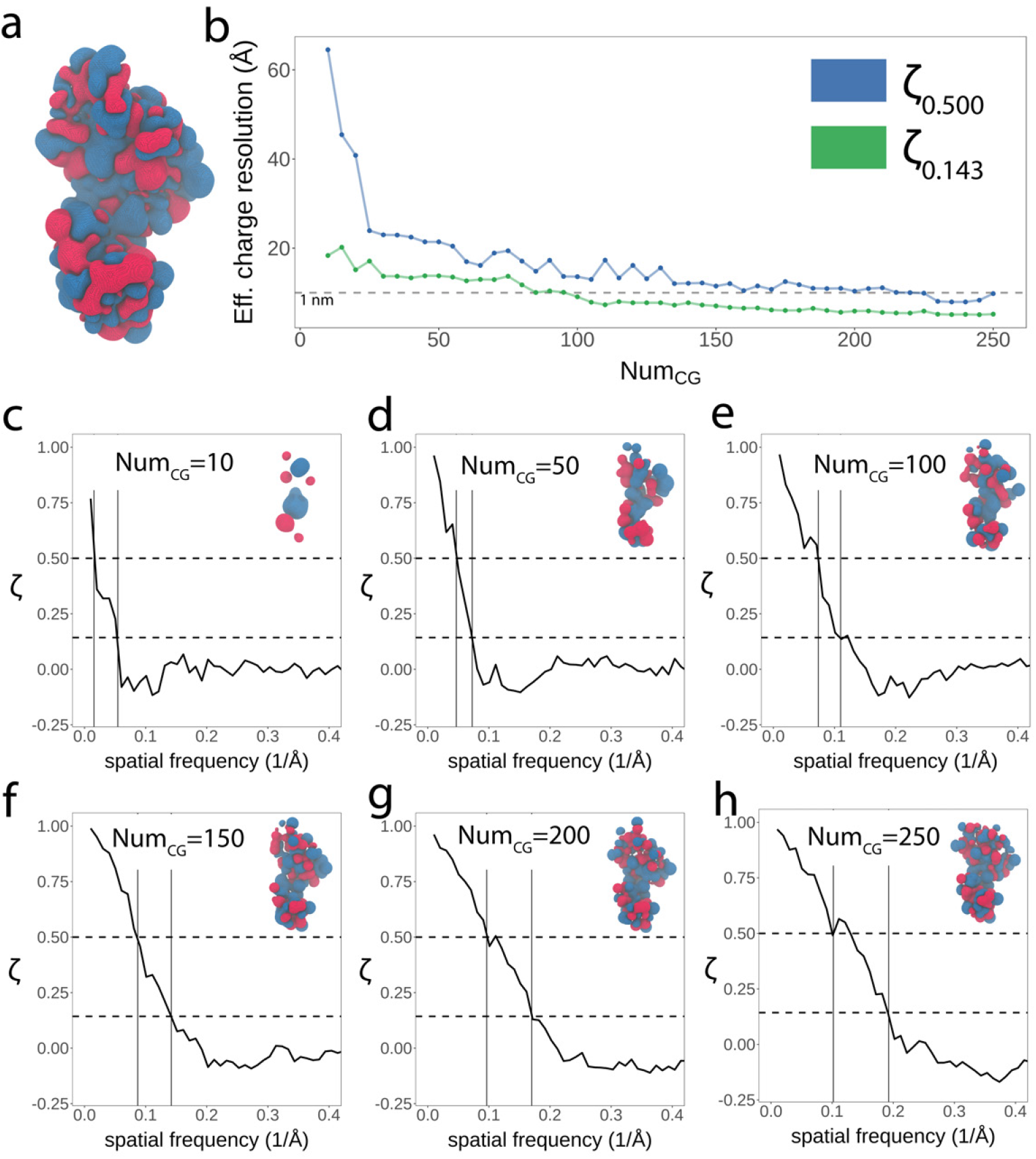
FSC analysis of CG granularity vs. effective charge density resolution. **a** Charge density of the all-atom reference structure of HIV-1 CA. Regions of positive and negative charge density are colored blue and red, respectively. **b** Effective charge density resolutions for models Num_CG_ ∈ [10, 250], plotted with two metrics: *ζ*_0.143_ and *ζ*_0.500_, green and blue, respectively. The dotted gray line represents a resolution of 1 nm. **c-h** Select charge density isosurfaces and FSC plots, Num_CG_ ∈ {10, 50, 100, 150, 200, and 250}. Charge densities are shown with identical isovalues and coloring to the density in panel a. Dashed horizontal lines denote each of the metrics employed, *ζ*_0.143_ and *ζ*_0.500_. Vertical solid lines in the plots denote the determination, via point of intersection, of effective resolution with *ζ*_0.143_ and *ζ*_0.500_ metrics.

**Figure S8:**
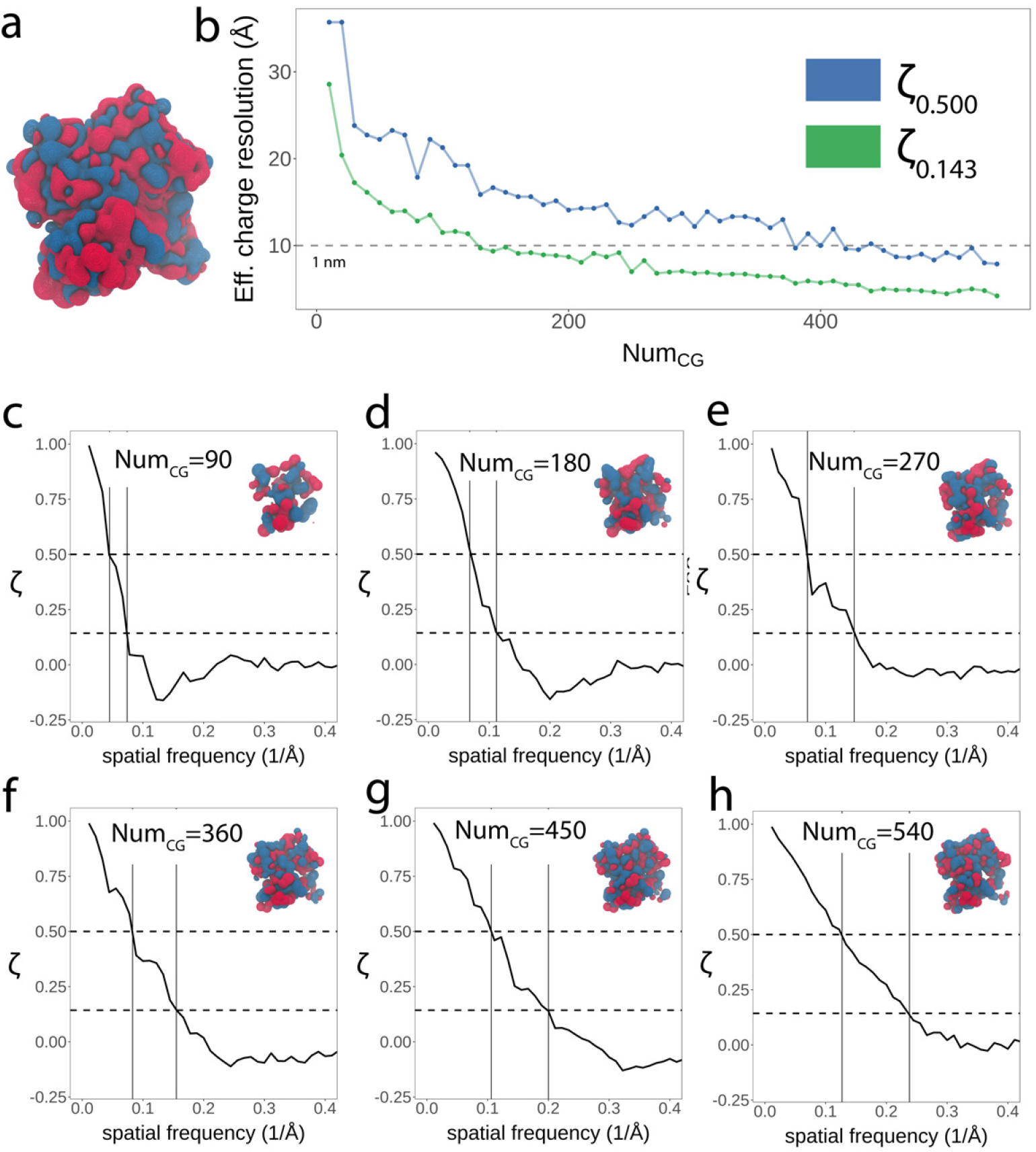
FSC analysis of CG granularity vs. effective charge density resolution. **a** Charge density of the all-atom reference structure of actin. Regions of positive and negative charge density are colored blue and red, respectively. **b** Effective charge density resolutions for models Num_CG_ ∈ [10, 540], plotted with two metrics: *ζ*_0.143_ and *ζ*_0.500_, green and blue, respectively. The dotted gray line represents a resolution of 1 nm. **c-h** Select charge density isosurfaces and FSC plots. Charge densities are shown with identical isovalues and coloring to the density in panel a. Dashed horizontal lines denote each of the metrics employed, *ζ*_0.143_ and *ζ*_0.500_. Vertical solid lines in the plots denote the determination, via point of intersection, of effective resolution with *ζ*_0.143_ and *ζ*_0.500_ metrics.

**Figure S9:**
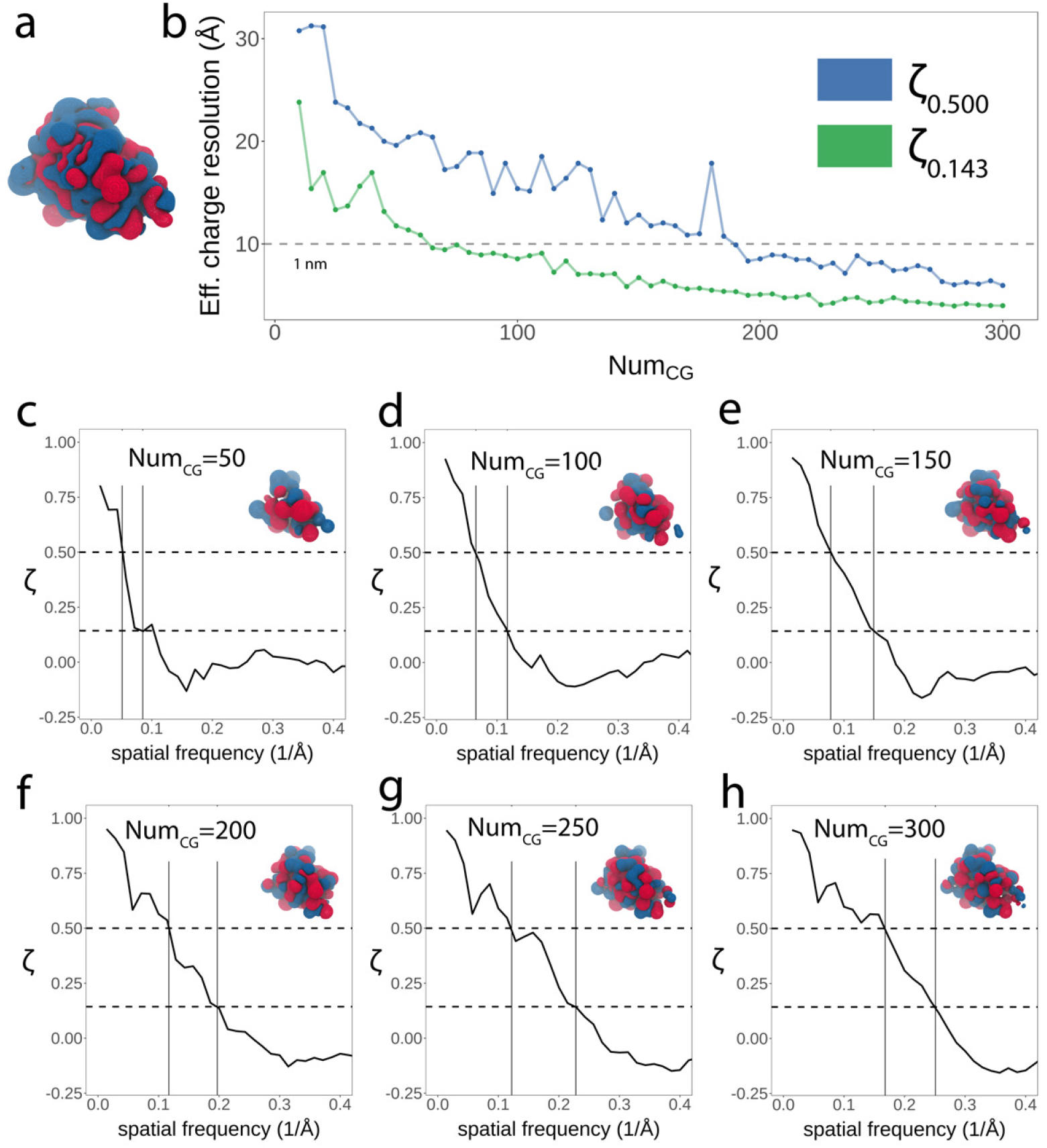
FSC analysis of CG granularity vs. effective charge density resolution. **a** Charge density of the all-atom reference structure of cofilin-2. Regions of positive and negative charge density are colored blue and red, respectively. **b** Effective charge density resolutions for models Num_CG_ ∈ [10, 300], plotted with two metrics: *ζ*_0.143_ and *ζ*_0.500_, green and blue, respectively. The dotted gray line represents a resolution of 1 nm. **c-h** Select charge density isosurfaces and FSC plots. Charge densities are shown with identical isovalues and coloring to the density in panel a. Dashed horizontal lines denote each of the metrics employed, *ζ*_0.143_ and *ζ*_0.500_. Vertical solid lines in the plots denote the determination, via point of intersection, of effective resolution with *ζ*_0.143_ and *ζ*_0.500_ metrics.

**Figure S10:**
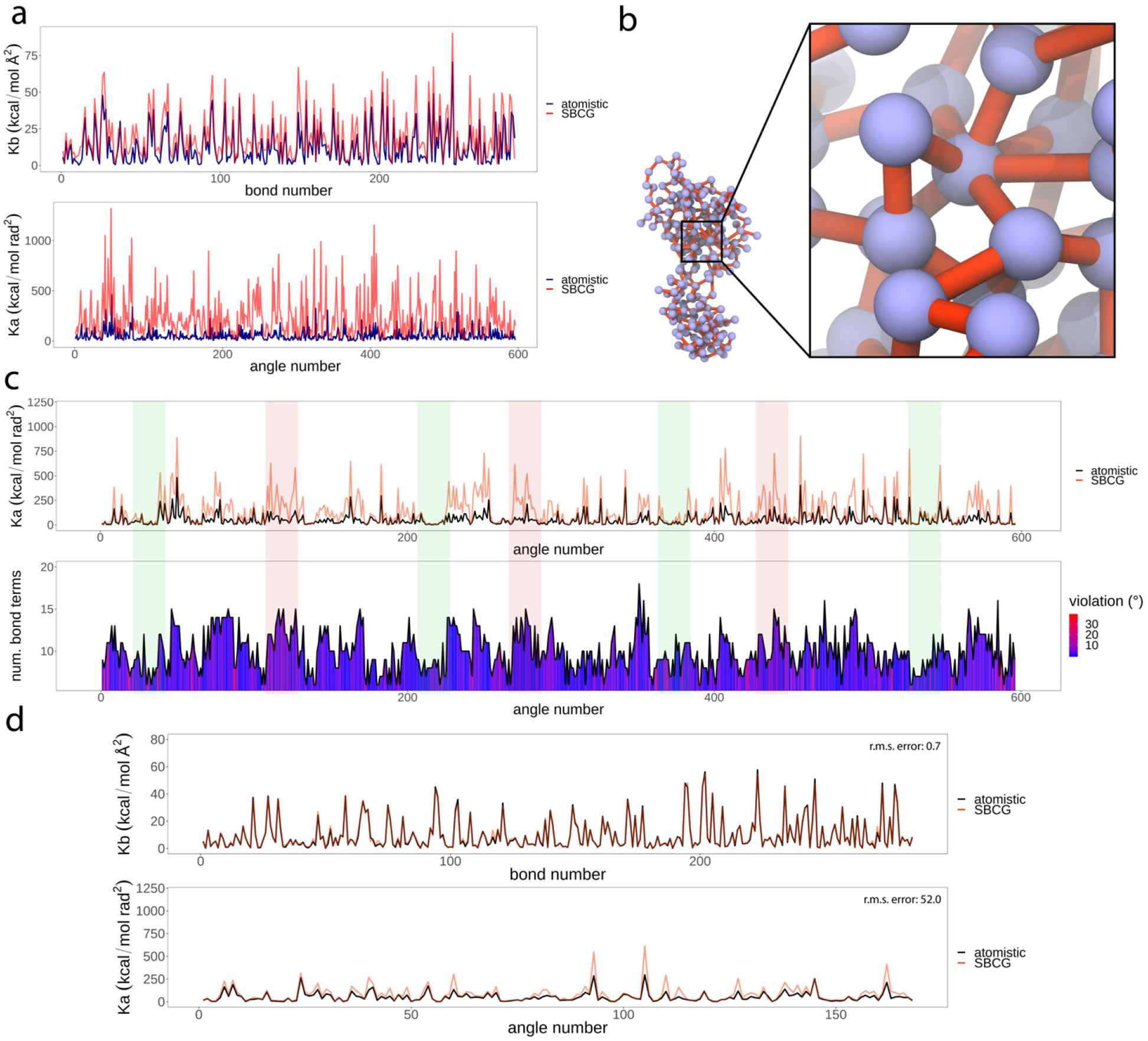
SBCG HIV-1 CA parameter optimization results. **a** Initial parameter set (red) derived from Boltzmann inversion of the all-atom trajectory (blue). Bond and angle (top and bottom, respectively) indices are given along the 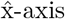, and the corresponding force constants are plotted along the 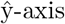. This plot demonstrates the poor quality of the initial fit, and motivates iterative refinement. **b** Example of a highly connected bead in the SBCG CA topology. **c** Analysis of angle parameters based on their bonded connectivity. The top panel shows angle parameters, converged to an unphysical state (orange), after many iterations of refinement (Movie M2). The bottom panel is aligned to the top and shows a trace of bonded connectivity, per angle parameter (for beads involved in the angle, all bonds connected to them are summed). The area under the latter trace is colored by violation, i.e., deviation of the angle value from the all-atom reference value. Regions of low connectivity correspond to good fit, shaded green. Regions of high connectivity correspond to bad fit, shaded red. **d** Converged parameter refinement following the pruning of redundant angle parameters. Final root-mean-square error (RMSE) values are shown.

**Figure S11:**
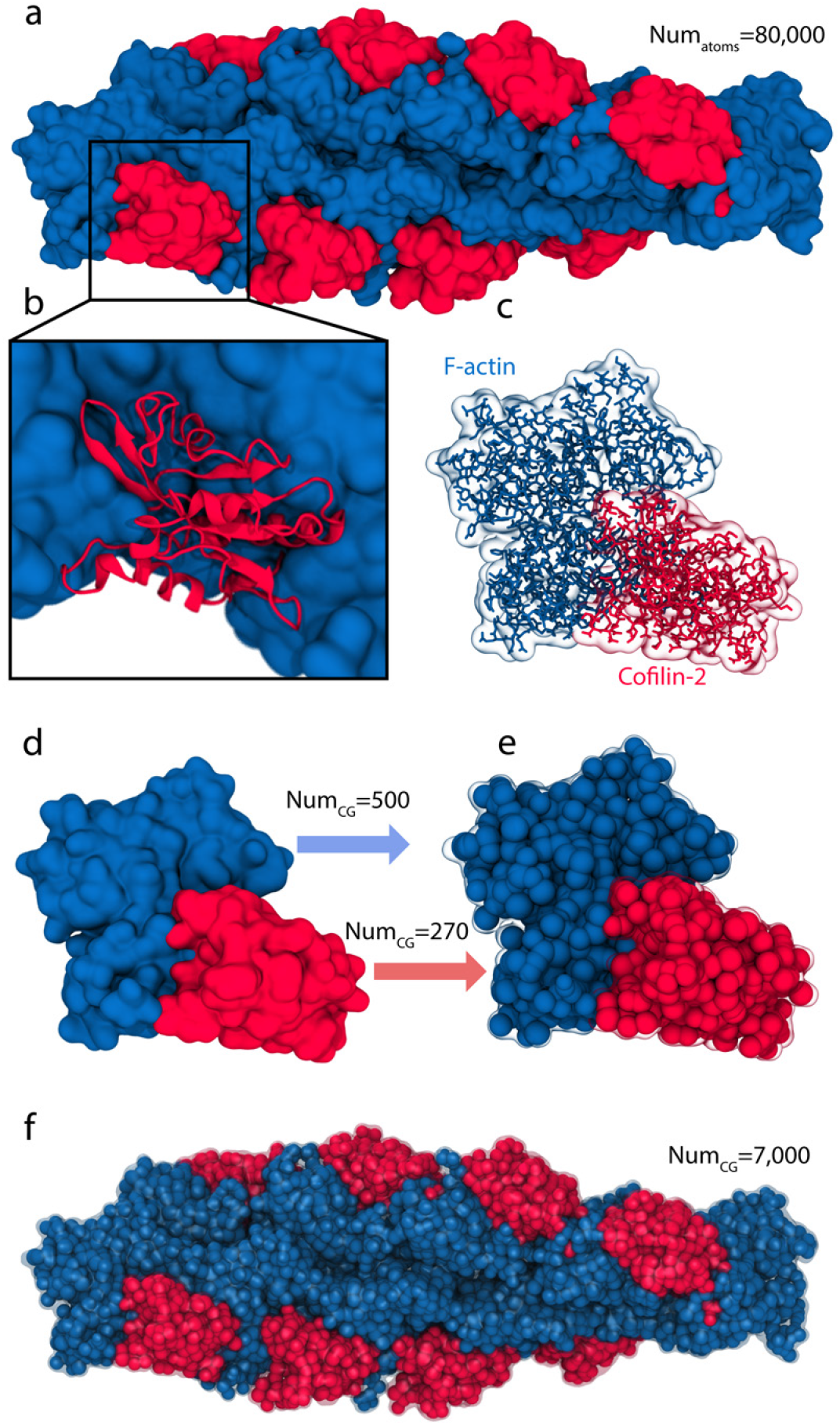
Heterogeneous multimeric assembly, cofilin-2 on actin filaments. **a** All-atom surface representation of the cofilin-2 and F-actin system. **b** Zoomed-in, secondary structure view of a cofilin-2 monomer in complex with actin, the latter shown in surface representation. **c** Atomistic view, without hydrogens, of each domain, cofilin-2 and F-actin, subjected to sub-nanometer shape based coarse graining. Molecular surfaces are shown transparently. **d,e** Atomistic surfaces, left, and the sub-nanometer SBCG models resulting from two independent calculations. Cofilin-2 modeling employed Num_CG_ = 270 and F-actin modeling employed Num_CG_ = 500, corresponding to approx. 11 atoms per bead for each protein domain. **f** Resulting multimeric SBCG cofilin-2 actin system after performing the mapping operation. The atomistic molecular surface is also shown transparently, demonstrating the quality of the resulting SBCG topology

**Figure M1:**
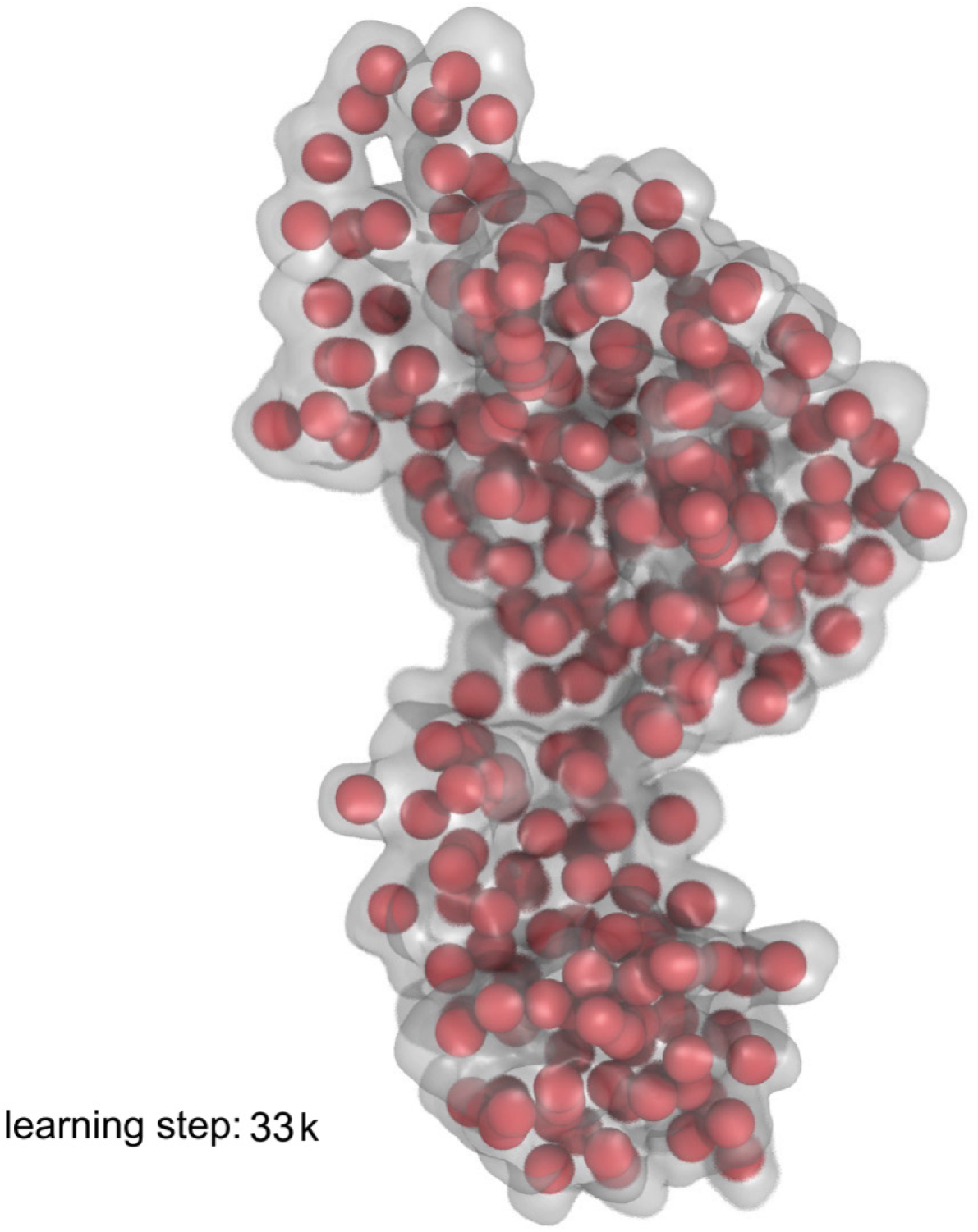
Animation of adaptation via the topology representing network.

**Figure M2:**
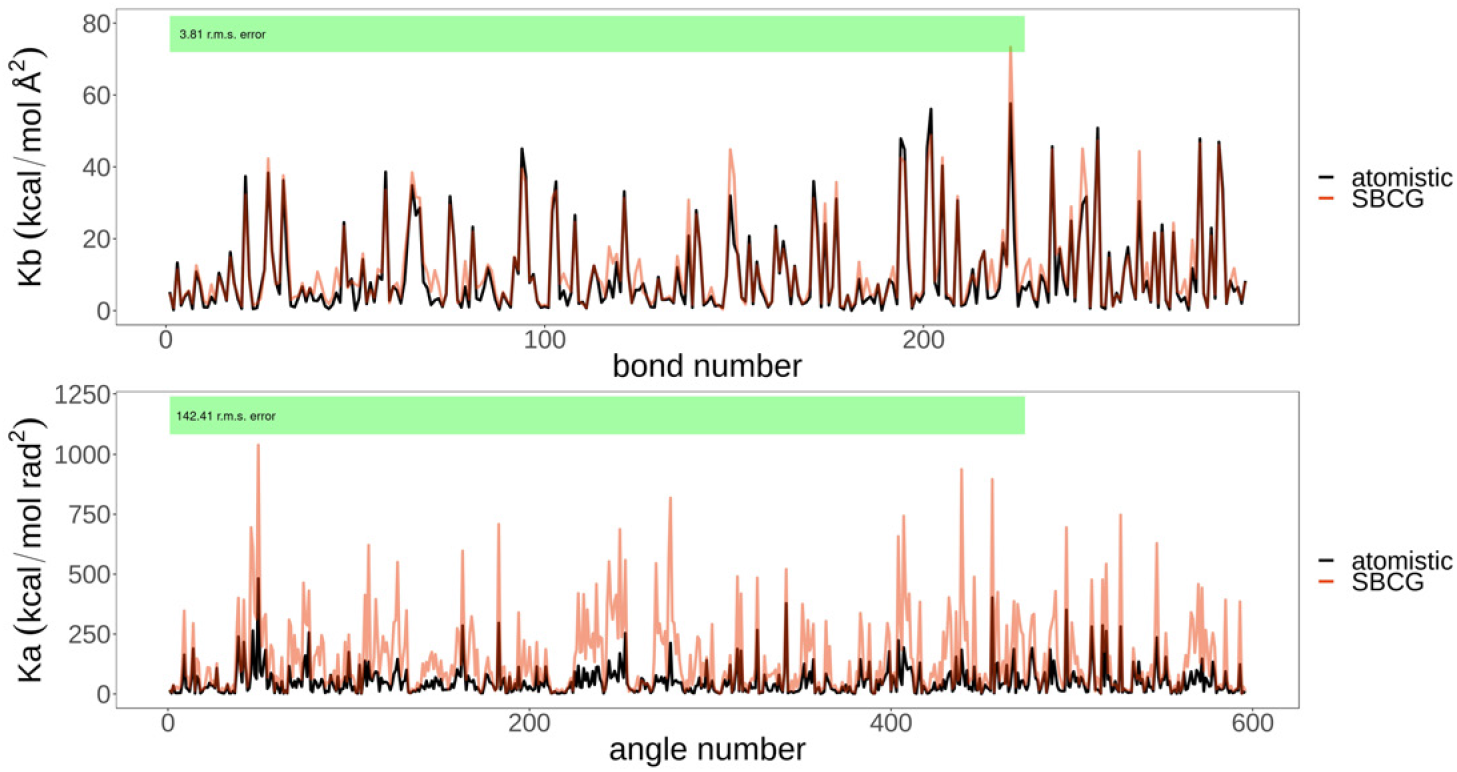
Animation of the iterative refinement protocol fit.

**Figure M3:**
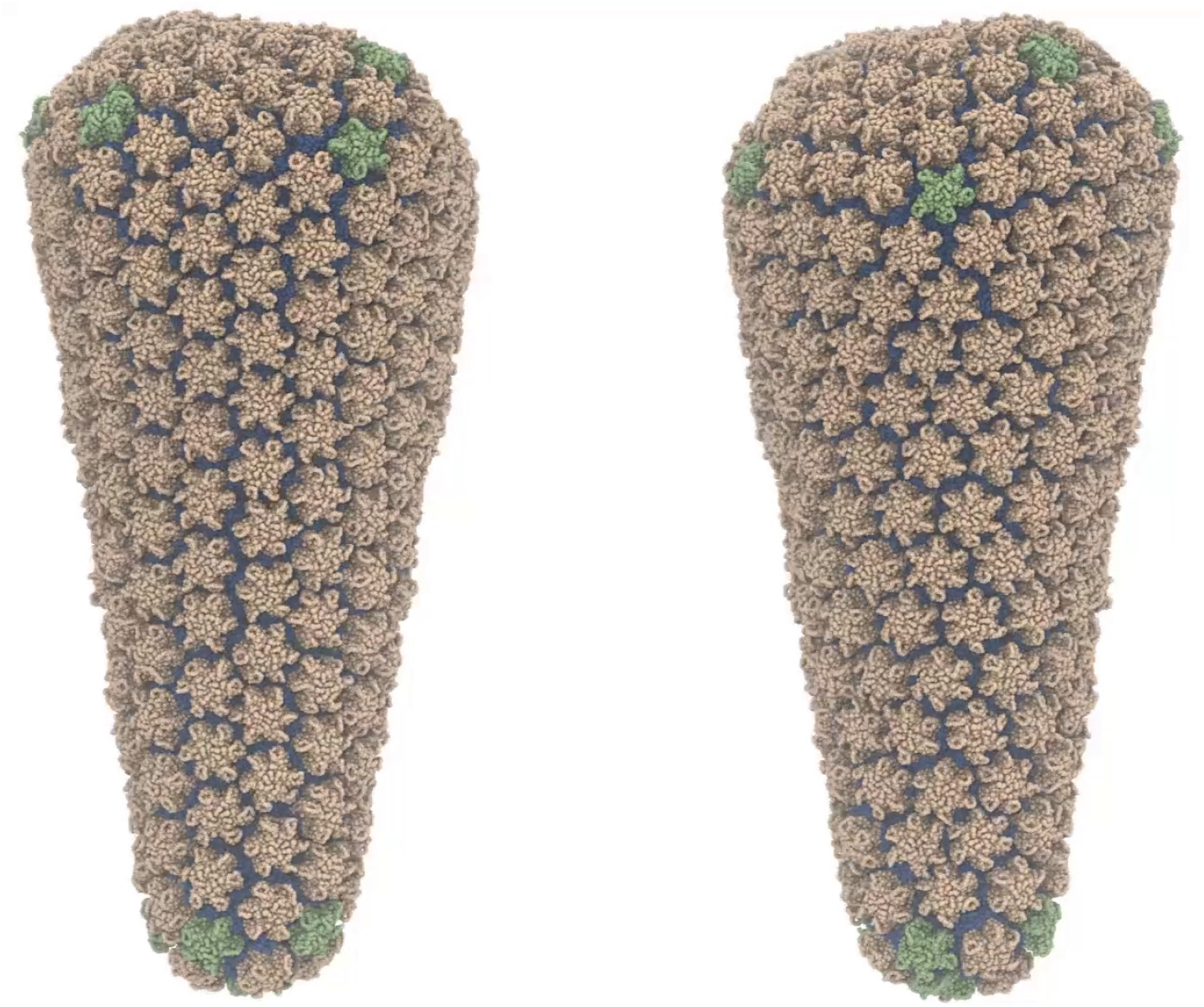
Movie of 225 ns of SBCG MD simulation of the HIV-1 conical capsid.

## Notes

### Competing Interest Statement

The authors have declared no competing interest.

## References

[1] Juan R Perilla, Boon Chong Goh, C Keith Cassidy, Bo Liu, Rafael C Bernardi, Till Rudack, Hang Yu, Zhe Wu, and Klaus Schulten. Molecular dynamics simulations of large macromolecular complexes. Current Opinion in Structural Biology, 31:64–74, 2015.

[2] David E Shaw, Martin M Deneroff, Ron O Dror, Jeffrey S Kuskin, Richard H Larson, John K Salmon, Cliff Young, Brannon Batson, Kevin J Bowers, Jack C Chao, et al. Anton, a special-purpose machine for molecular dynamics simulation. Communications of the ACM, 51(7):91–97, 2008.

[3] Michael Levitt and Arieh Warshel. Computer simulation of protein folding. Nature, 253(5494):694–698, 1975.

[4] Steve O Nielsen, Carlos F Lopez, Goundla Srinivas, and Michael L Klein. Coarse grain models and the computer simulation of soft materials. Journal of Physics: Condensed Matter, 16(15):R481, 2004.

[5] Thomas A Knotts IV, Nitin Rathore, David C Schwartz, and Juan J De Pablo. A coarse grain model for DNA. The Journal of Chemical Physics, 126(8):02B611, 2007.

[6] Marc Baaden and Siewert J Marrink. Coarse-grain modelling of protein–protein interactions. Current Opinion in Structural Biology, 23(6):878–886, 2013.

[7] Alisa S Wolberg, Leping Li, Wing-Fai Cheung, Nobuko Hamaguchi, Lee G Pedersen, and Darrel W Stafford. Characterization of *γ*-carboxyglutamic acid residue 21 of human factor IX. Biochemistry, 35(32):10321–10327, 1996.

[8] Mikio Sakai, Minami Abe, Yusuke Shigeto, Shin Mizutani, Hiroyuki Takahashi, Axelle Viré, James R Percival, Jiansheng Xiang, and Christopher C Pain. Verification and validation of a coarse grain model of the DEM in a bubbling fluidized bed. Chemical Engineering Journal, 244:33–43, 2014.

[9] Siewert J Marrink, H Jelger Risselada, Serge Yefimov, D Peter Tieleman, and Alex H De Vries. The MARTINI force field: coarse grained model for biomolecular simulations. The Journal of Physical Chemistry B, 111(27):7812–7824, 2007.

[10] Luca Monticelli, Senthil K Kandasamy, Xavier Periole, Ronald G Larson, D Peter Tieleman, and Siewert-Jan Marrink. The MARTINI coarse-grained force field: extension to proteins. Journal of Chemical Theory and Computation, 4(5):819–834, 2008.

[11] Xavier Periole and Siewert-Jan Marrink. The MARTINI coarse-grained force field. Biomolecular Simulations, pages 533–565, 2013.

[12] James F Dama, Anton V Sinitskiy, Martin McCullagh, Jonathan Weare, Benoît Roux, Aaron R Dinner, and Gregory A Voth. The theory of ultra-coarse-graining. 1. General principles. Journal of Chemical Theory and Computation, 9(5):2466–2480, 2013.

[13] Michael F Hagan and Roya Zandi. Recent advances in coarse-grained modeling of virus assembly. Current Opinion in Virology, 18:36–43, 2016.

[14] Farzaneh Mohajerani, Evan Sayer, Christopher Neil, Koe Inlow, and Michael F Hagan. Mechanisms of scaffold-mediated microcompartment assembly and size control. ACS Nano, 15(3):4197–4212, 2021.

[15] Alvin Yu, Katarzyna A Skorupka, Alexander J Pak, Barbie K Ganser-Pornillos, Owen Pornillos, and Gregory A Voth. *TRIM5α* self-assembly and compartmentalization of the HIV-1 viral capsid. Nature Communications, 11(1):1–10, 2020.

[16] Alvin Yu, Elizabeth MY Lee, John AG Briggs, Barbie K Ganser-Pornillos, Owen Pornillos, and Gregory A Voth. Strain and rupture of HIV-1 capsids during uncoating. Proceedings of the National Academy of Sciences, 119(10):e2117781119, 2022.

[17] Tamar Schlick, Eric Barth, and Margaret Mandziuk. Biomolecular dynamics at long timesteps: Bridging the timescale gap between simulation and experimentation. Annual Review of Biophysics and Biomolecular Structure, 26(1):181–222, 1997.

[18] Raghunath Chelakkot, Arvind Gopinath, Lakshminarayanan Mahadevan, and Michael F Hagan. Flagellar dynamics of a connected chain of active, polar, Brownian particles. Journal of The Royal Society Interface, 11(92):20130884, 2014.

[19] Hongmei Jian, Alexander V Vologodskii, and Tamar Schlick. A combined wormlike-chain and bead model for dynamic simulations of long linear DNA. Journal of Computational Physics, 136(1):168–179, 1997.

[20] William George Noid, Jhih-Wei Chu, Gary S Ayton, Vinod Krishna, Sergei Izvekov, Gregory A Voth, Avisek Das, and Hans C Andersen. The multiscale coarse-graining method. I. A rigorous bridge between atomistic and coarse-grained models. The Journal of Chemical Physics, 128(24):244114, 2008.

[21] Martín Soñora, Leandro Martinez, Sergio Pantano, and Matías R Machado. Wrapping up viruses at multiscale resolution: optimizing PACKMOL and SIRAH execution for simulating the zika virus. Journal of Chemical Information and Modeling, 61(1):408–422, 2021.

[22] Wei Han, Cheuk-Kin Wan, Fan Jiang, and Yun-Dong Wu. PACE force field for protein simulations. 1. Full parameterization of version 1 and verification. Journal of Chemical Theory and Computation, 6(11):3373–3389, 2010.

[23] Wei Han, Cheuk-Kin Wan, and Yun-Dong Wu. PACE force field for protein simulations. 2. Folding simulations of peptides. Journal of Chemical Theory and Computation, 6(11):3390–3402, 2010.

[24] Jiang Wang, Simon Olsson, Christoph Wehmeyer, Adri’a Pérez, Nicholas E Charron, Gianni De Fabritiis, Frank Noé, and Cecilia Clementi. Machine learning of coarse-grained molecular dynamics force fields. ACS Central Science, 5(5):755–767, 2019.

[25] Henry Chan, Mathew J Cherukara, Badri Narayanan, Troy D Loeffler, Chris Benmore, Stephen K Gray, and Subramanian KRS Sankaranarayanan. Machine learning coarse grained models for water. Nature Communications, 10(1):1–14, 2019.

[26] Aleksander EP Durumeric and Gregory A Voth. Adversarial-residual-coarse-graining: Applying machine learning theory to systematic molecular coarse-graining. The Journal of Chemical Physics, 151(12):124110, 2019.

[27] James L McDonagh, Ardita Shkurti, David J Bray, Richard L Anderson, and Edward O Pyzer-Knapp. Utilizing machine learning for efficient parameterization of coarse grained molecular force fields. Journal of Chemical Information and Modeling, 59(10):4278–4288, 2019.

[28] Brooke E Husic, Nicholas E Charron, Dominik Lemm, Jiang Wang, Adri’a Pérez, Maciej Majewski, Andreas Krämer, Yaoyi Chen, Simon Olsson, Gianni de Fabritiis, et al. Coarse graining molecular dynamics with graph neural networks. The Journal of Chemical Physics, 153(19):194101, 2020.

[29] Yuwei Zhang, Yunchu Wang, Fei Xia, Zexing Cao, and Xin Xu. Accurate and efficient estimation of Lennard–Jones interactions for coarse-grained particles via a potential matching method. Journal of Chemical Theory and Computation, 2022.

[30] Beste Bayramoglu and Roland Faller. Coarse-grained modeling of polystyrene in various environments by iterative Boltzmann inversion. Macromolecules, 45(22):9205–9219, 2012.

[31] Dirk Reith, Mathias Pütz, and Florian Müller-Plathe. Deriving effective mesoscale potentials from atomistic simulations. Journal of Computational Chemistry, 24(13):1624–1636, 2003.

[32] Sergei Izvekov and Gregory A Voth. A multiscale coarse-graining method for biomolecular systems. The Journal of Physical Chemistry B, 109(7):2469–2473, 2005.

[33] Aviel Chaimovich and M Scott Shell. Coarse-graining errors and numerical optimization using a relative entropy framework. The Journal of Chemical Physics, 134(9):094112, 2011.

[34] Zhiyong Zhang, Jim Pfaendtner, Andrea Grafmüller, and Gregory A Voth. Defining coarse-grained representations of large biomolecules and biomolecular complexes from elastic network models. Biophysical Journal, 97(8):2327–2337, 2009.

[35] Marissa G Saunders and Gregory A Voth. Coarse-graining of multiprotein assemblies. Current Opinion in Structural Biology, 22(2):144–150, 2012.

[36] Marissa G Saunders and Gregory A Voth. Coarse-graining methods for computational biology. Annual Review of Biophysics, 42:73–93, 2013.

[37] Jodi Kraus, Ryan W Russell, Elena Kudryashova, Chaoyi Xu, Nidhi Katyal, Juan R Perilla, Dmitri S Kudryashov, and Tatyana Polenova. Magic angle spinning NMR structure of human cofilin-2 assembled on actin filaments reveals isoform-specific conformation and binding mode. Nature Communications, 13(1):1–12, 2022.

[38] James C Phillips, David J Hardy, Julio DC Maia, John E Stone, João V Ribeiro, Rafael C Bernardi, Ronak Buch, Giacomo Fiorin, Jéôme Hénin, Wei Jiang, et al. Scalable molecular dynamics on CPU and GPU architectures with NAMD. The Journal of Chemical Physics, 153(4):044130, 2020.

[39] Robert A Dick, Kaneil K Zadrozny, Chaoyi Xu, Florian KM Schur, Terri D Lyddon, Clifton L Ricana, Jonathan M Wagner, Juan R Perilla, Barbie K Ganser-Pornillos, Marc C Johnson, et al. Inositol phosphates are assembly co-factors for HIV-1. Nature, 560(7719):509–512, 2018.

[40] Tao Ni, Yanan Zhu, Zhengyi Yang, Chaoyi Xu, Yuriy Chaban, Tanya Nesterova, Jiying Ning, Till Böcking, Michael W Parker, Christina Monnie, et al. Structure of native HIV-1 cores and their interactions with IP6 and CypA. Science Advances, 7(47):eabj5715, 2021.

[41] Juan R Perilla and Klaus Schulten. Physical properties of the HIV-1 capsid from all-atom molecular dynamics simulations. Nature Communications, 8(1):1–10, 2017.

[42] Thomas Martinetz and Klaus Schulten. Topology representing networks. Neural Networks, 7(3):507–522, 1994.

[43] Anton Arkhipov, Peter L Freddolino, and Klaus Schulten. Stability and dynamics of virus capsids described by coarse-grained modeling. Structure, 14(12):1767–1777, 2006.

[44] Anton Arkhipov, Ying Yin, and Klaus Schulten. Four-scale description of membrane sculpting by BAR domains. Biophysical Journal, 95(6):2806–2821, 2008.

[45] Anton Arkhipov, Wouter H Roos, Gijs JL Wuite, and Klaus Schulten. Elucidating the mechanism behind irreversible deformation of viral capsids. Biophysical Journal, 97(7):2061–2069, 2009.

[46] Anton Arkhipov, Ying Yin, and Klaus Schulten. Membrane-bending mechanism of amphiphysin N-BAR domains. Biophysical Journal, 97(10):2727–2735, 2009.

[47] Hang Yu and Klaus Schulten. Membrane sculpting by F-BAR domains studied by molecular dynamics simulations. PLoS Computational Biology, 9(1):e1002892, 2013.

[48] Chris Rycroft. Voro++: A three-dimensional Voronoi cell library in C++. Technical report, Lawrence Berkeley National Lab.(LBNL), Berkeley, CA (United States), 2009.

[49] POV-Ray. Persistence of Vision Raytracer version 3.7.0. Persistence of Vision Pty. Ltd., 2004.

[50] D.O. Hebb. The Organization of Behavior: A Neuropsychological Theory. Taylor & Francis Group, 2002.

[51] Tony Lindeberg. A computational theory of visual receptive fields. Biological cybernetics, 107(6):589–635, 2013.

[52] Thomas M Martinetz, Stanislav G Berkovich, and Klaus J Schulten. ‘Neural-gas’ network for vector quantization and its application to time-series prediction. IEEE Transactions on Neural Networks, 4(4):558–569, 1993.

[53] William Humphrey, Andrew Dalke, and Klaus Schulten. VMD: visual molecular dynamics. Journal of Molecular Graphics, 14(1):33–38, 1996.

[54] George Harauz and Marin van Heel. Exact filters for general geometry three dimensional reconstruction. Optik., 73(4):146–156, 1986.

[55] Pawel A Penczek. Resolution measures in molecular electron microscopy. In Methods in Enzymology, volume 482, pages 73–100. Elsevier, 2010.

[56] W O Saxton and W Baumeister. The correlation averaging of a regularly arranged bacterial cell envelope protein. Journal of Microscopy, 127(2):127–138, 1982.

[57] Guang Tang, Liwei Peng, Philip R Baldwin, Deepinder S Mann, Wen Jiang, Ian Rees, and Steven J Ludtke. EMAN2: an extensible image processing suite for electron microscopy. Journal of Structural Biology, 157(1):38–46, 2007.

[58] NVIDIA. cuFFT library version 11.7.0. NVIDIA Corp., 2022.

[59] Richard Henderson, Andrej Sali, Matthew L Baker, Bridget Carragher, Batsal Devkota, Kenneth H Downing, Edward H Egelman, Zukang Feng, Joachim Frank, Nikolaus Grigorieff, et al. Outcome of the first electron microscopy validation task force meeting. Structure, 20(2):205–214, 2012.

[60] Scott M Stagg, Alex J Noble, Michael Spilman, and Michael S Chapman. ResLog plots as an empirical metric of the quality of cryo-EM reconstructions. Journal of Structural Biology, 185(3):418–426, 2014.

[61] Peter B Rosenthal and Richard Henderson. Optimal determination of particle orientation, absolute hand, and contrast loss in single-particle electron cryomicroscopy. Journal of Molecular Biology, 333(4):721–745, 2003.

[62] Sjors HW Scheres and Shaoxia Chen. Prevention of overfitting in cryo-EM structure determination. Nature Methods, 9(9):853–854, 2012.

[63] Hossein Ali Karimi-Varzaneh, Hu-Jun Qian, Xiaoyu Chen, Paola Carbone, and Florian Müller-Plathe. IBIsCO: A molecular dynamics simulation package for coarse-grained simulation. Journal of Computational Chemistry, 32(7):1475–1487, 2011.

[64] Martin Hanke. Well-posedness of the iterative Boltzmann inversion. Journal of Statistical Physics, 170(3):536–553, 2018.

[65] Chaoyi Xu, Douglas K Fischer, Sanela Rankovic, Wen Li, Robert A Dick, Brent Runge, Roman Zadorozhnyi, Jinwoo Ahn, Christopher Aiken, Tatyana Polenova, et al. Permeability of the HIV-1 capsid to metabolites modulates viral DNA synthesis. PLoS Biology, 18(12):e3001015, 2020.

[66] Juan R Perilla, Oliver Beckstein, Elizabeth J Denning, and Thomas B Woolf. Computing ensembles of transitions from stable states: Dynamic importance sampling. Journal of Computational Chemistry, 32(2):196–209, 2011.

[67] James C Phillips, Rosemary Braun, Wei Wang, James Gumbart, Emad Tajkhorshid, Elizabeth Villa, Christophe Chipot, Robert D Skeel, Laxmikant Kale, and Klaus Schulten. Scalable molecular dynamics with NAMD. Journal of Computational Chemistry, 26(16):1781–1802, 2005.

[68] Daan Frenkel and Berend Smit. Understanding molecular simulation: from algorithms to applications, volume 1. Elsevier, 2001.

[69] Peter E Kloeden and Eckhard Platen. Stochastic differential equations. In Numerical solution of stochastic differential equations, pages 103–160. Springer, 1992.

[70] Florian Sammüller and Matthias Schmidt. Adaptive Brownian dynamics. The Journal of Chemical Physics, 155(13):134107, 2021.

[71] Tom Darden, Darrin York, and Lee Pedersen. Particle mesh Ewald: An N log (N) method for Ewald sums in large systems. The Journal of Chemical Physics, 98(12):10089–10092, 1993.

[72] Ulrich Essmann, Lalith Perera, Max L Berkowitz, Tom Darden, Hsing Lee, and Lee G Pedersen. A smooth particle mesh Ewald method. The Journal of Chemical Physics, 103(19):8577–8593, 1995.

[73] John E Stone, David J Hardy, Ivan S Ufimtsev, and Klaus Schulten. GPU-accelerated molecular modeling coming of age. Journal of Molecular Graphics and Modelling, 29(2):116–125, 2010.

